# Three-dimensional Interrogation of Cell Types and Instinctive Behavior in the Periaqueductal Gray

**DOI:** 10.1101/2022.06.27.497769

**Authors:** Eric Vaughn, Stephen Eichhorn, Won Jung, Xiaowei Zhuang, Catherine Dulac

## Abstract

The periaqueductal gray (PAG) is a critical midbrain hub that relays information from the forebrain to motor and autonomic brainstem centers to orchestrate instinctive behaviors. The current organization of the PAG into four main radial columns lacks the resolution needed to account for the vast range of PAG functions. Using spatially resolved single-cell transcriptomic measurements, we uncovered widespread transcriptional heterogeneity in the PAG with >100 excitatory and inhibitory neuronal populations, which further assemble into 19 spatial metaclusters. We explored the transcriptional and spatial logic of PAG function during instinctive behaviors and demonstrated the regional recruitment of cell types for distinct behaviors. Unexpectedly, certain behaviors trigger differential spatial activation patterns within given cell types, illustrating the complexity of PAG molecular and functional 3D organization. The newly uncovered spatial motifs and high precision cellular map of instinctive behavior in the PAG open new avenues for a mechanistic understanding of PAG function.

## Introduction

In order to survive and reproduce, animals must exhibit an array of instinctive behavioral and physiological responses such as fighting, mating, escaping, and parenting, which involve coordinated somatic and autonomic functions. The periaqueductal gray matter (PAG) is an evolutionarily conserved midbrain structure that orchestrates these instinctive functions by integrating cortical, subcortical, and spinal inputs before conveying this information to downstream motor and autonomic pathways (Carrive, 1993). Despite its broad involvement in behavior control, the precise molecular and cellular organization of the PAG that underlies its various functions has remained elusive. Decades of neuro- histochemical and tracing analyses have established a radial subdivision of the PAG into dorsomedial, dorsolateral, lateral, and ventrolateral columns (dm, dl, l, vl-PAG), each characterized by distinct inputs, projections, and molecular profiles (Bandler and Keay, 1991; Carrive, 1993). At the functional level, the dorsal PAG is thought to be mainly involved in active coping behaviors, such as flight, mating and aggression, while the ventral PAG regulates more passive coping functions such as freezing, nursing and sleeping. However, recent studies have pointed to a more complex organization in which distinct radial columns participate in multiple and overlapping functions. For instance, the lPAG has been shown to commonly mediate vocalization (Tschida et al., 2019), attack (Falkner et al., 2020), defense (Wang et al., 2019), mating (Han et al., 2017; Yamada and Kawata, 2014), predation (Han et al., 2017), and itch (Gao et al., 2019), while a specific behavior such as fear jointly recruits dlPAG, lPAG, and vlPAG. Such widespread and intermingled activity highlights the need for a more comprehensive and high resolution understanding of the spatial and molecular organization underlying PAG functions (Falkner et al., 2020; Rossier et al., 2021; Tschida et al., 2019; Zhong et al., 2019).

The PAG lacks anatomically identifiable subnuclei, such as those defined in amygdaloid and hypothalamic areas. As a result, experiments assessing specific PAG functions have largely relied on available columnar and inhibitory versus excitatory subdivisions. However, these broad delineations fail to adequately parcellate the PAG into molecularly and functionally relevant units (Esteban Masferrer et al., 2020; Falkner et al., 2020; Reis et al., 2021; Tovote et al., 2016). Recent advances in single-cell and spatial transcriptomics provide a unique opportunity to more precisely dissect the spatial, molecular and cell type-specific organization underlying distinct behavioral and homeostatic functions in the PAG. Using a combination of single nucleus RNA-sequencing (snRNA-seq) (Lake et al., 2016) and Multiplexed Error-Robust Fluorescence in Situ Hybridization (MERFISH) (Moffitt et al., 2016), we have molecularly and functionally defined neuronal clusters in the PAG and surrounding regions. We identified 144 subpopulations of neurons and, using their three-dimensional spatial attributes, grouped them into 19 metaclusters sharing the same spatial motifs. Using a panel of immediate early genes (IEGs), we characterized the involvement of each cell cluster and spatial metacluster in a variety of behaviors in males and females, such as adult and infant-mediated aggression, parenting, mating, predator exposure and exercise. We find clusters displaying behavior-specific (i.e., just predator exposure) and behaviorally shared (i.e., predator exposure and parenting) activation patterns that may reflect the merging of complex responses into functional outputs relevant to multiple behaviors.

Remarkably, we find that behavior-induced activity can be further spatially distributed *within* a given cluster, thus uncovering a parcellation of PAG functions according to both cell type and spatial coordinates. Lastly, using transsynaptic anterograde and retrograde viral tracing we further link molecularly defined PAG populations to input from functionally defined upstream nuclei. Our work provides a novel framework for examining the organization and function of spatially related cell populations and unveiled the organization and logic of a brain region critical for instinctive and homeostatic responses.

## Results

### Transcriptomic classification of cell types in the PAG

To identify transcriptionally distinct cell populations within the PAG, we dissected the PAG and surrounding structures from the posterior thalamus to the anterior cerebellum (∼2.5 by 2.5 by 3 mm, Bregma -2.5 to -5.5 mm) and performed snRNA-Seq on ∼100,000 cells (Figure 1A; Methods). After quality control and removal of non-PAG neuronal populations, we clustered the data and used this information to select a 262-gene MERFISH library (Table 1) based on the following criteria: 1) filter differentially expressed genes expressed in less than 40% of all neurons to focus on transcripts that can best delineate clusters; 2) employ a minimum redundancy maximum relevancy algorithm to select a panel of the ∼200 most informative genes necessary to distinguish clusters; 3) select differentially expressed genes of specific biological relevance not already included in 1 & 2; 4) Train a radial support vector machine classifier on this gene set to classify snRNA-seq cells, iteratively removing and adding genes to provide each cluster with high prediction accuracy (Fig. S1 A-D; Methods). We biased this approach to maximally assess diversity in neuronal populations, while selecting a minimum set of genes distinguishing non-neuronal cells. Additionally, we added 26 IEGs to define cellular activity during a given behavior (Table 1).

**Figure 1:**
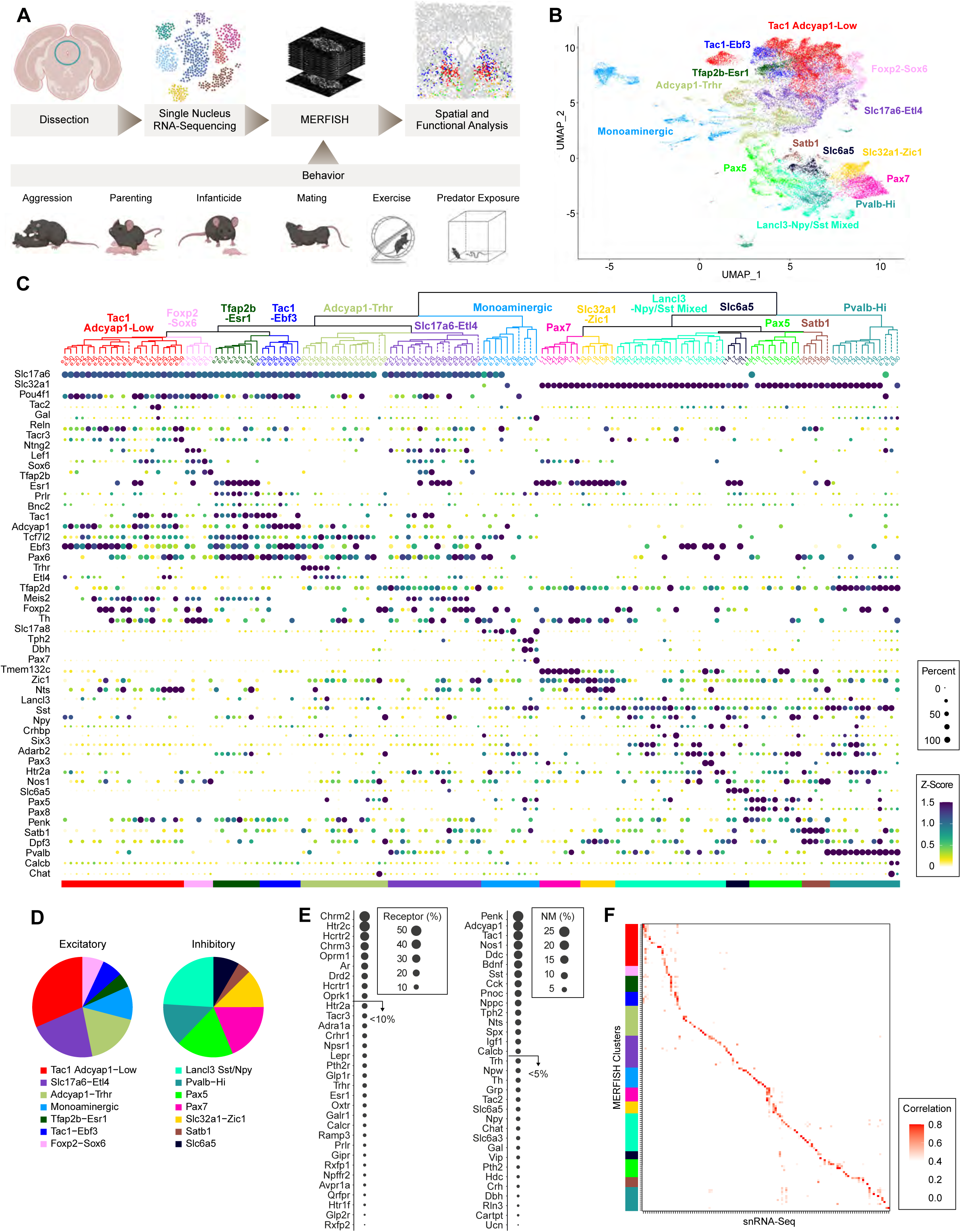
Molecular categorization of PAG MERFISH Clusters. **A**: Overview of experimental paradigm. **B**: UMAP representation where colors represent 14 hierarchically defined transcriptomic groups. **C**: Dendrogram representing the hierarchical organization of cluster based on gene expression; color schema identical as in B. Dotted branches on dendrogram represent clusters not classically defined as part of the PAG. Dot size within matrix represents the percent of cells expressing a given gene listed on the y-axis; color of dot represents mean cluster z-score for a given gene. **D**: Piechart distribution depicting the relative abundance of cells within each transcriptomic metacluster for excitatory (left) or inhibitory (right) subsets. **E**: Percent of cells expressing specific neuromodulatory (top) or neuromodulatory receptor (bottom) genes. **F**: MERFISH and snRNA-Seq Pearson cluster correlation matrix.

**Table 1.**

| **Genes** | **IEGs** |
| --- | --- |
| **Gpc5** | Nrn1 |
| **Adarb2** | Homer1 |
| **Nxph1** | Rheb |
| **Etl4** | Nr4a3 |
| **Kcnq5** | Bdnf |
| **Il1rapl2** | Jund |
| **Rgs6** | Myc |
| **Htr2c** | Rgs2 |
| **Sorcs1** | Nptx2 |
| **Ptprt** | Plk2 |
| **Chrm2** | Nr4a1 |
| **Zfpm2** | Junb |
| **Gpc6** | Plat |
| **Slit3** | Fos |
| **Tph2** | Jun |
| **Meis2** | Egr1 |
| **Gria3** | Npas4 |
| **Ntng1** | Fosl1 |
| **Ebf1** | Inhba |
| **Cacna2d3** | Fosl2 |
| **Rnf220** | Arc |
| **Cacna2d1** | Fosb |
| **Ddc** | Ptgs2 |
| **Sema6d** | Egr4 |
| **Chrm3** | Egr2 |
| **Tenm3** | Egr3 |
| **Hs3st4** |  |
| **Col25a1** |  |
| **Robo1** |  |
| **Dach1** |  |
| **Arpp21** |  |
| **Sox6** |  |
| **Pbx3** |  |
| **Slc18a2** |  |
| **Nos1** |  |
| **Tcf4** |  |
| **Kcnh8** |  |
| **Ebf2** |  |
| **Nfib** |  |
| **Megf11** |  |
| **Fign** |  |
| **Tox** |  |
| **Zeb2** |  |
| **Tmem132d** |  |
| **Foxp2** |  |
| **Cadps2** |  |
| **Sv2c** |  |
| **Cep112** |  |
| **Ptprk** |  |
| **Nfia** |  |
| **Zfhx3** |  |
| **Syt6** |  |
| **Zfhx4** |  |
| **Rorb** |  |
| **Chat** |  |
| **Pde1a** |  |
| **Tmem132c** |  |
| **Ndst4** |  |
| **Gad2** |  |
| **Pde7b** |  |
| **Tshz2** |  |
| **Otx2os1** |  |
| **Calcr** |  |
| **Dscaml1** |  |
| **Epha4** |  |
| **Rln3** |  |
| **Pde3a** |  |
| **Tcf7l2** |  |
| **Satb1** |  |
| **Reln** |  |
| **Pdzd2** |  |
| **Pou6f2** |  |
| **Syt2** |  |
| **Ndst3** |  |
| **Tll1** |  |
| **Col11a1** |  |
| **Adgrv1** |  |
| **Lrmda** |  |
| **Bnc2** |  |
| **Dpf3** |  |
| **Sox5** |  |
| **Dbh** |  |
| **Fras1** |  |
| **Cdh4** |  |
| **X9030622O22Rik** |  |
| **Meis1** |  |
| **Ankfn1** |  |
| **Zfp536** |  |
| **Vwa5b1** |  |
| **4-Sep** |  |
| **Mgll** |  |
| **Sgpp2** |  |
| **Cck** |  |
| **Zfp521** |  |
| **Maf** |  |
| **Necab1** |  |
| **Gfra1** |  |
| **Bcl11a** |  |
| **Cdh23** |  |
| **Pax3** |  |
| **Tgfbr2** |  |
| **Mitf** |  |
| **Htr1f** |  |
| **Th** |  |
| **Angpt1** |  |
| **Cobll1** |  |
| **Sema3e** |  |
| **Adra1a** |  |
| **Penk** |  |
| **Sema3d** |  |
| **Ebf3** |  |
| **Stac2** |  |
| **Sst** |  |
| **Npsr1** |  |
| **Slc17a8** |  |
| **Vip** |  |
| **Lef1** |  |
| **Slc17a6** |  |
| **Spp1** |  |
| **Dlk1** |  |
| **Pde1c** |  |
| **B230323A14Rik** |  |
| **Cnih3** |  |
| **Pappa** |  |
| **Baiap3** |  |
| **Tac1** |  |
| **Oprm1** |  |
| **Hcrtr2** |  |
| **St18** |  |
| **Lhx4** |  |
| **Bcl11b** |  |
| **Fam163a** |  |
| **Sema3a** |  |
| **Hcrtr1** |  |
| **Chst9** |  |
| **Gm38505** |  |
| **Gal** |  |
| **Sec61b** |  |
| **Slc6a3** |  |
| **Ntng2** |  |
| **Lmx1b** |  |
| **Pax5** |  |
| **Ntn1** |  |
| **Tfap2d** |  |
| **Nnat** |  |
| **Sim1** |  |
| **Col19a1** |  |
| **Calcb** |  |
| **Gata3** |  |
| **Spon1** |  |
| **Tanc1** |  |
| **Htr2a** |  |
| **Prlr** |  |
| **Esrrb** |  |
| **Tfap2b** |  |
| **Rnf207** |  |
| **Qrfpr** |  |
| **Fstl1** |  |
| **Igf1** |  |
| **Ar** |  |
| **Npy** |  |
| **Nts** |  |
| **Isl1** |  |
| **Gli3** |  |
| **Rab3b** |  |
| **Fbn2** |  |
| **Ttc6** |  |
| **Oprk1** |  |
| **Esr1** |  |
| **C1ql1** |  |
| **Slc32a1** |  |
| **Prox1** |  |
| **Glis3** |  |
| **Iqgap2** |  |
| **Adamts19** |  |
| **Lancl3** |  |
| **Sema3c** |  |
| **Six3** |  |
| **Col23a1** |  |
| **Mecom** |  |
| **Wls** |  |
| **Drd2** |  |
| **Rxfp1** |  |
| **Etfdh** |  |
| **Osbpl3** |  |
| **Gpr139** |  |
| **Slc6a5** |  |
| **En1** |  |
| **Foxo1** |  |
| **Gpc3** |  |
| **Nr4a2** |  |
| **Nrip3** |  |
| **Lepr** |  |
| **Crhr1** |  |
| **Tacr3** |  |
| **Rab37** |  |
| **Spx** |  |
| **Gipr** |  |
| **C1ql2** |  |
| **Pax8** |  |
| **Glp2r** |  |
| **Pvalb** |  |
| **Pnoc** |  |
| **Pax7** |  |
| **Pou4f1** |  |
| **Arhgef17** |  |
| **Sall3** |  |
| **Chrna6** |  |
| **Crhbp** |  |
| **Syt10** |  |
| **Fxyd6** |  |
| **Cers2** |  |
| **Cfl2** |  |
| **Tmem39b** |  |
| **Trh** |  |
| **Npffr2** |  |
| **Zic1** |  |
| **Trhr** |  |
| **Lmo1** |  |
| **Rxfp2** |  |
| **X1700011I03Rik** |  |
| **Pax6** |  |
| **Pth2r** |  |
| **C1ql3** |  |
| **Bmp6** |  |
| **Plscr4** |  |
| **Col24a1** |  |
| **Gad1os** |  |
| **Synpo2** |  |
| **Ttr** |  |
| **Chchd10** |  |
| **Crh** |  |
| **Surf1** |  |
| **Nkx2.2** |  |
| **Galr1** |  |
| **Grp** |  |
| **Nppc** |  |
| **Adcyap1** |  |
| **Ramp3** |  |
| **Mtnr1a** |  |
| **Mrc1** |  |
| **Npw** |  |
| **Agt** |  |
| **Hdc** |  |
| **Avpr1a** |  |
| **Slc17a7** |  |
| **Glp1r** |  |
| **Tac2** |  |
| **Oxtr** |  |
| **Vtn** |  |
| **Pdgfra** |  |
| **X1500015O10Rik** |  |
| **Pth2** |  |
| **Lum** |  |
| **Arhgap45** |  |
| **Slc1a3** |  |
| **Cldn5** |  |
| **Mog** |  |
| **Snap25** |  |
| **Hepacam** |  |
| **Ucn** |  |
| **Cartpt** |  |

Using the gene set defined above, we performed MERFISH on coronal PAG slices (2 x 2 x 2.5 mm, Bregma -2.7 to 5.2 mm) across 31 replicates covering animals of both sexes and various behavior conditions. From the ∼2.5 million cells profiled, graph-based clustering defined 144 neuronal and 10 non-neuronal clusters. To join neuronal profiles into similar transcriptomic groups, we hierarchically sorted the average cluster expression and applied a dynamic tree cut to parse clusters into 14 groups (Fig. 1B and C). These groups were divided into excitatory or inhibitory branches where, numerically, excitatory neurons outnumbered inhibitory neurons 71.4% to 28.6%. A majority (83.2%) of neurons expressed at least one neuropeptide, where Tac1 and Adcyap1 were particularly widespread and variable in their expression, but selective to excitatory neurons. Many excitatory clusters were defined by specific expression of Adcyap1, Tac1, Cck, Esr1, Trhr, Grp, Gal, and Galr1, with rare populations expressing Crh, Npsr1, Cartpt, Ramp3, or Prlr (Fig. 1C-E; Fig. S1E). By comparison, many inhibitory clusters were defined by Pnoc, Hctr1, Npy, Htr2a, and Adra1a, with rare populations expressing Trh, Qrfpr, or Htr1f. A subset of inhibitory neurons also expressed the glycine transporter Slc6a5. Other neuromodulatory markers such as Penk, Sst, Nos1, Nts, Igf1, Oxtr, and Tacr3 were less specific to inhibitory or excitatory clusters. Expression of the sex hormone receptors Ar and Pgr was widespread (Fig. S1F and S1G), while expression of Esr1 and Prlr was more specific and often colocalized— particularly in a population of Tfap2b-Esr1 neurons (Fig. 1C). Interestingly, ependymal and radial glia-like cells lining the aqueduct expressed region-specific transcription factors (ex. Zic1 and Pax7), as well as membrane trafficking proteins (synaptotagmin-10 (Syt10)), neuropeptides (Pdyn) and neuropeptide receptors (Npsr1, Oxtr, Qrfpr, Ednrb) (Fig. S1H-O), potentially endowing them with signaling abilities via the ventricular system, as previously suggested (Veening and Barendregt, 2010).

Additionally, we profiled several PAG-encompassed or surrounding structures, including the nucleus of Darkschewitsch, oculomotor, trochlear, Edinger-Westphal, laterodorsal tegmental nuclei (LDT), and dorsal raphe (DR). However, because these structures were not the focus of this study, genes were not included in the MERFISH library to thoroughly parse the corresponding cell types. Nevertheless, we captured a wide variety of monoaminergic and cholinergic neurons expressing Th, Ddc, Tph2, Dbh, Slc6a3, Slc18a2, and Chat, comprising the serotonergic DR; cholinergic oculomotor, trochlear, and LDT; as well as dopaminergic vlPAG and sparse noradrenergic neurons in caudal PAG (caudal PAG).

Comparison between neuronal snRNA-seq and MERFISH cluster expression profiles showed a mean max Pearson correlation of 0.68, demonstrating a high degree of correspondence across technologies (Fig. 1F) with MERFISH adding increased resolution in many cases due to higher cell count and sensitivity to peptide and receptor gene expression that are particularly sparse in snRNA-seq. Thus, our data defines a vast transcriptomic array of PAG cell populations expressing a wide variety of neuromodulatory genes, consistent with the role of the PAG in a myriad of functions.

### Spatial characterization of PAG cell types

We next sought to characterize the spatial representation of cell populations across the whole PAG. We first assessed the distribution of major cell classes and found that most nonneuronal cells were evenly dispersed, except for ependymal cells and a population of radial glia-like cells appearing near, but not limited to the circumventricular subcommissural organ (Fig. 2A; Fig. S2A). Inhibitory and excitatory neurons were largely intermingled, with higher density of inhibitory populations in the rostral PAG, and lower density in the caudal half of the lPAG (Fig. 2B; Fig. S2A-C). An average ∼4mm2 coronal slice contained ∼35 abundantly represented neuronal clusters (>30 neurons in a respective slice), suggesting a high diversity in neuronal populations at any given level of the PAG structure (Fig. S2D). To determine the extent to which profiles of gene expression are determined by space, we plotted the three-dimensional coordinates onto the uniform manifold approximation projection (UMAP), which was built using only gene expression. For nonneuronal cell classes, spatial values appeared to be randomly distributed within each class’s UMAP space, except for a slight dorsoventral gradient in astrocytes (Fig. 2 C-F). However, for neurons, a clear relationship emerged between transcriptional profiles and spatial coordinates, particularly in the dorsoventral and rostrocaudal axes. Excitatory neurons displayed a clear gradient from dorsal to ventral in UMAP space, whereas inhibitory neurons largely fell into separate dorsal or ventral groups, as evidenced by fewer intermediate cells in Fig. 2E.

**Figure 2:**
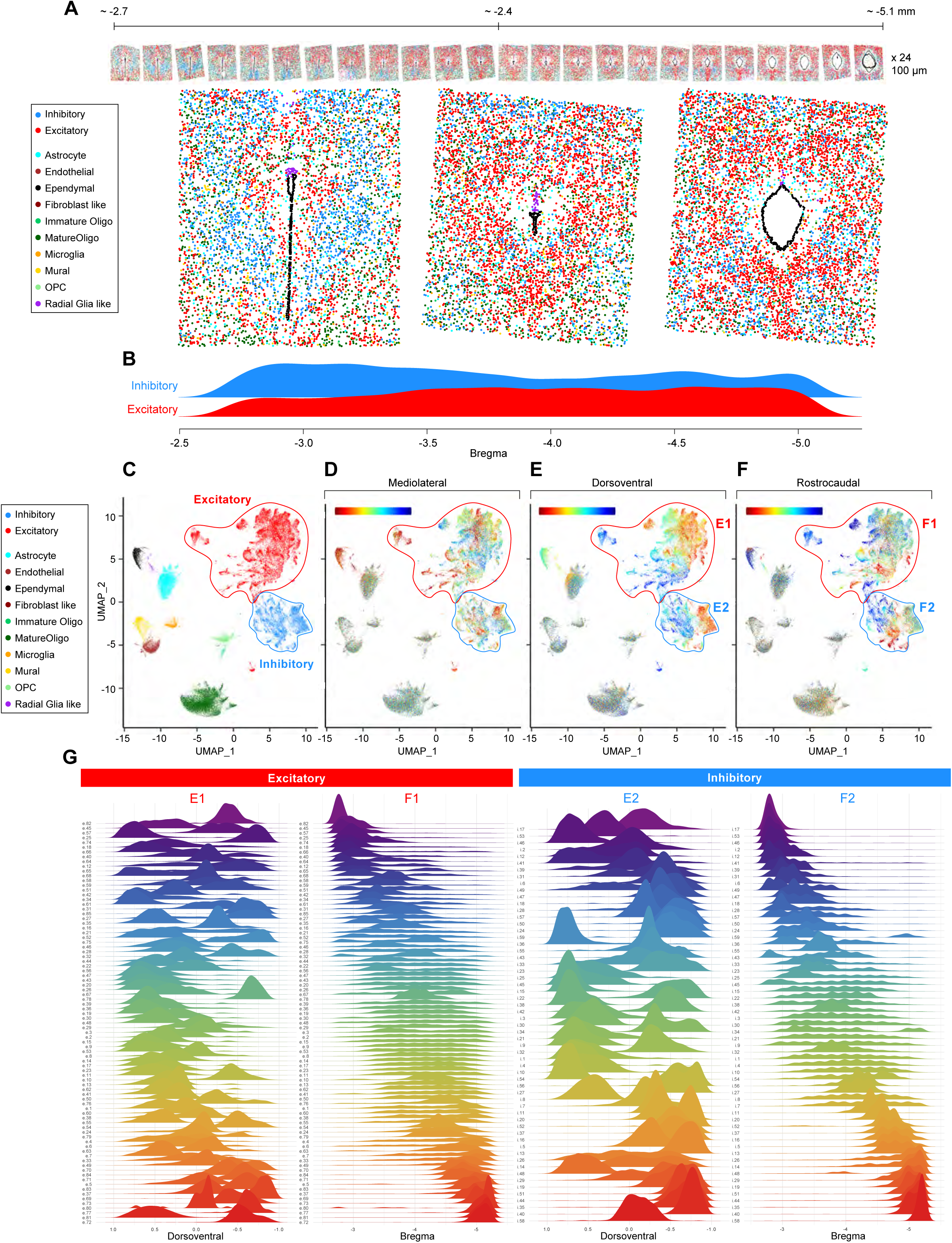
Spatial Distribution of Major Types and Neuronal Clusters. **A**: Coronal MERFISH sections showing the distribution of major cell classes across the rostral (left) to caudal (right) length of the PAG. **B**: Density plot showing the distribution of Inhibitory or Excitatory neurons across the rostrocaudal axis. **C-F**: From left to right: representation of major cell types in UMAP space, followed by color-coded mediolateral, dorsoventral, and rostrocaudal location of cells plotted on the same UMAP. **G**: Neuronal distribution of each excitatory and inhibitory cluster along the rostrocaudal (left) and dorsoventral (right) axes.

Electrophysiological and opto/chemogenetic stimulations have established a delineation of various behaviors and autonomic functions along the rostrocaudal axis of the PAG (Bandler and Depaulis, 1991; Sukikara et al., 2006; Tovote et al., 2016; Tschida et al., 2019; Zhong et al., 2019). Consistent with this organization, subsets of neuronal clusters were only present in specific locations along the rostrocaudal axis (Fig. 2G). By contrast, other clusters were present along nearly the entire length of the PAG, thus contributing to the classically described columnar organization of the PAG (Discussed further below). Excitatory clusters extended typically longer along the rostrocaudal axis (mean SD ± .48mm) and therefore tended to be more evenly dispersed, while inhibitory clusters tended to extend shorter distances along the Bregma axis (mean SD ± .39mm, *P* < 0.0005). Of the 85 excitatory clusters, 48 had a ± SD length over 1mm, while only 16/59 inhibitory clusters were longer than 1mm. The presence of discrete inhibitory populations along the rostrocaudal axis could endow them with the ability to uniquely modulate local circuits along this axis. Indeed, previous work has demonstrated the critical roles of local inhibitory interneurons to subserve specific functions within the PAG such as freezing (Tovote et al., 2016), sleep (Zhong et al., 2019), and ultrasound vocalization (Chen et al., 2021; Tovote et al., 2016). These findings provide a new framework for understanding how site-specific activation of the PAG may trigger a large repertoire of distinct behavioral and autonomic responses.

### Defining Three-Dimensional Spatial Metaclusters

The PAG has classically been subdivided into four longitudinal columns (i.e. dm-, dl-, l-, and vlPAG). However, due to the lack of anatomically distinct sub-nuclei and the so far limited in situ transcriptional measurements in the PAG, the precise molecular and spatial organization of neuronal types in the PAG—and the extent to which they obey the proposed columnar organization—are yet to be defined. The precise spatial distributions of individual cell types obtained from MERFISH measurements allowed us to directly address this question and offered an opportunity to identify previously unknown spatial motifs as well as define the neuronal populations that share these motifs. To define which neuronal populations shared similar 3D topologies, we used spatial information for all neuronal types to build a nearest-neighbor graph for each neuron, i.e., computing the cell type composition of the nearest neighbors for each neuron. We then performed hierarchical clustering analysis to identify spatial metaclusters (SMCs), within which neurons share similar neighborhood composition (Fig. 3A; Methods). This analysis yielded 19 SMCs, providing both a robust molecular basis for the classical columnar PAG subdivisions, as well as considerably refining and revising to this organization (Fig. S3).

**Figure 3:**
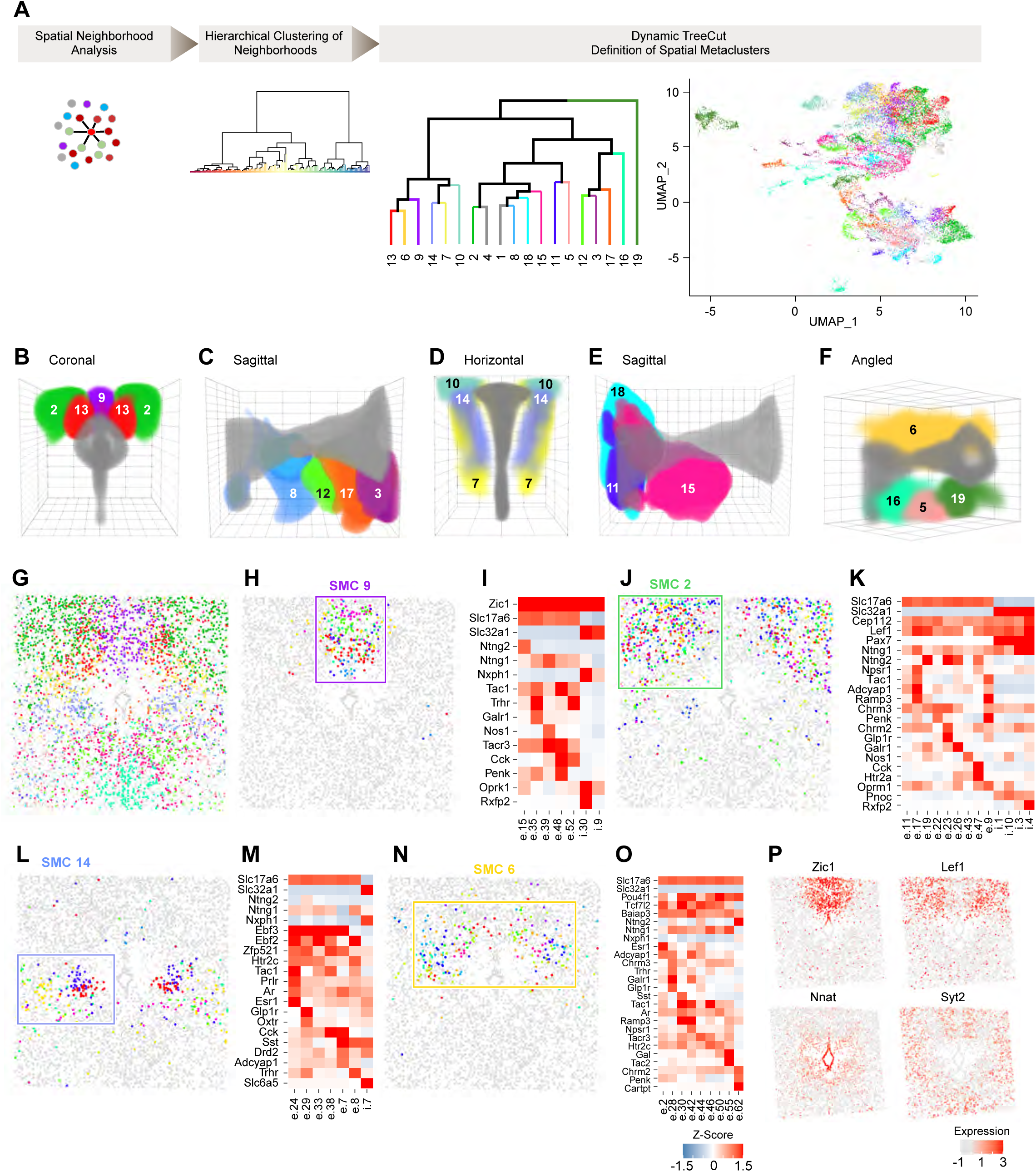
Definition and Molecular Characterization of Spatial Metaclusters. A: Overview of spatial network analysis: Dynamic TreeCut definition of Spatial Metaclusters colored by each metacluster and plotted on representative coronal sections to illustrate location. UMAP colored by metacluster. B: Three-dimensional coronal view of SMCs 9, 13, and 2 (purple, red, and green, respectively). C: Three-dimensional sagittal view of SMCs 8, 12, 17, and 3, and 2 (blue, green, orange, and, purple, respectively). D: Three-dimensional sagittal view of SMCs 18, 11, and 15 (cyan, blue, and pink, respectively) E: Three-dimensional horizontal view of SMCs 7, 14, and 10 (blue, yellow, teal, respectively). F: Angled Three-dimensional view of SMCs 6, 5, 16, and 19 (gold, light green, pink, dark green, respectively) G: Coronal section detailing SMCs plotted at -4.0 Bregma, colored as in B-F. H: Illustration of clusters (colored by cluster) within dorsomedial SMC 9 (purple in G). Gray cells represent cells outside of SMC of interest. I: Z-score expression heatmap of clusters within F. J-O: Same as H and I but for SMCs 2, 14, and 6, respectively. P: Coronal slices showing z-score expression of Zic1, Lef1 (top), Nnat and Syt2 (bottom).

Refining columnar subdivisions, we found that SMC 9 corresponded to dmPAG; SMCs 2 and 13, respectively corresponded to the outer and inner regions of dlPAG (Fig. 3B). The vlPAG was broken into four separate SMCs that were present at progressively more caudal positions, respectively (8, 12, 17, and 3; Fig. 3C). SMCs 7, 14, and 10 respectively corresponded to the outer, inner, and caudal lPAG (Fig. 3D). We also defined several SMCs confined to rostral PAG, identifying them as spatially and molecularly separate (11, 18, 15, Fig. 3E). We also define SMCs confined to the rostral ventromedial portion of the PAG, the oculomotor and supraoculomotor-related nuclei, and dorsal raphe (16, 5, and 19, respectively, Fig. 3F). As an exception to columnar organization, SMC 6 contained nine clusters that formed a dorsal arch starting from lPAG, before gradually becoming confined to dlPAG at ∼-4.8 AP (Fig. 3F; Fig. S3). We also noted that some SMCs, while initially confined to separate columns in the rostral zones, collide in caudal zones: dmPAG SMC 9 and dlPAG SMC 13 were cleanly distinguished at -3.8 AP yet were largely intermixed in the dorsomedial region of caudal PAG (Fig. S3). Hence, specific clusters may be confined to one classically defined PAG column, extend to several previously defined columns, or transition through a given column to another one. Additionally, PAG columns can be subdivided radially by inner and outer regions, or by rostral or caudal segments.

Altogether, these data uncover a novel framework for the precise identification of distinct modes of organization (both columnar and non-columnar modes), and identify a high level of cellular heterogeneity within each spatially defined PAG domain.

To confirm the robustness of our SMCs, we separately performed spatial clustering by directly comparing the spatial distributions of individual clusters, instead of the neighborhood compositions of individual clusters as described above. These two analyses yielded nearly identical SMCs (Fig. S4A). As a negative control, we performed SMC analysis on simulated cluster distributions where the cell number and physical dimensions of each neuronal cluster were kept the same as the experimentally derived values, but the cluster centroid positions were randomly placed in the PAG (Methods).

Compared to real SMCs, SMCs derived from such randomized neuronal clusters showed much less spatial confinement (Fig. S4B, C; p < 10-5), confirming the existence of spatial commonalities between multiple clusters that assemble into specific SMCs.

Transcriptionally, clusters forming a given SMC shared expression motifs that defined their spatiality, while simultaneously displaying strong internal heterogeneities. For example, the dmPAG SMC 9 was defined by the common expression of Zic1, while the outer dlPAG SMC 2 expressed Cep112, Lef1, and Pax7 in inhibitory neurons (Fig.3 H-K; Fig. S4D). However, within SMCs, distinct arrays of neuropeptides, neuromodulatory receptors, and cell signaling molecules were expressed in different cell clusters, revealing a surprising diversity in neuromodulatory signaling in what are often treated as homogenous columns of cells (Fig. 3H-O). For instance, within SMC 9, Cck, Nos1, Penk, Oprk1, Galr1, and Rxfp2 were differentially expressed between clusters, with similar observations made in other SMCs. Hence, SMCs provide a new delineation for molecularly and spatially defined and separable neuronal populations within the PAG, while also highlighting the internal complexity within these populations that may serve as the basis for their computations.

To further assess the spatial molecular landscape of the PAG, we tested genes for spatial autocorrelation using Moran’s i (*i*; Methods) and uncovered spatially heterogeneous gene expression patterns. A total of 54 genes had an *i* > 0.20, suggesting that a variety of partially spatially overlapping genes contribute to the definition of SMCs (Table 2; *p* < 0.01). Accordingly, while genes like Zic1 and Lef1 were relatively confined to specific SMCs, many genes with spatially confined expression patterns were expressed across several SMCs (Fig. 3P). For example, Nnat and Syt2 demarcate radial distance from the aqueduct (Fig. 3P). Thus, while decades of research have subdivided the PAG into coarse columnar subdivisions, our analysis highlights a substantially more complex organization, and more broadly, provides a new framework for investigating spatial motifs of molecularly defined cell types.

**Table 2.**
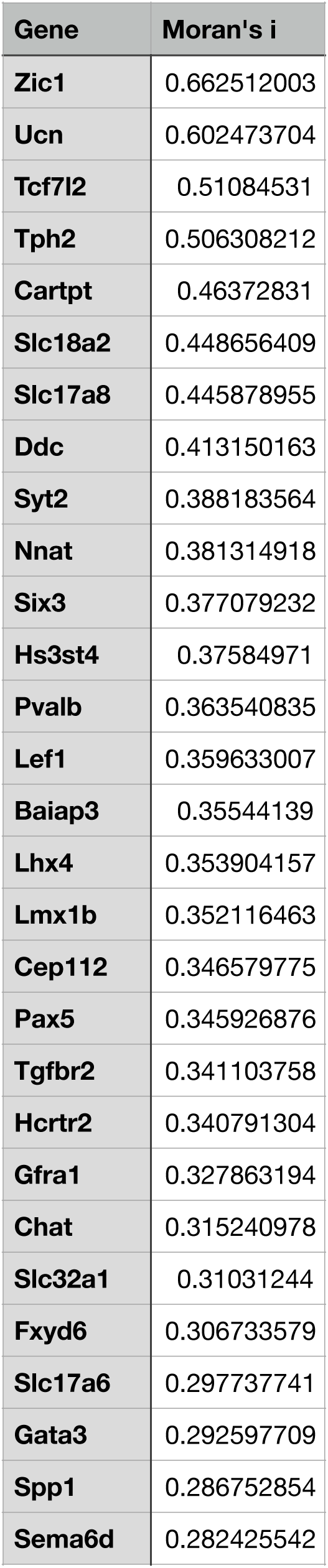
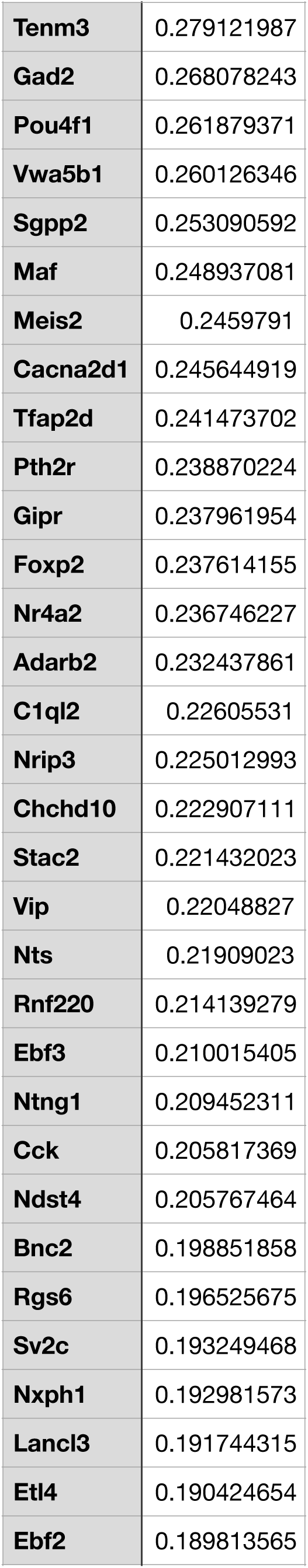
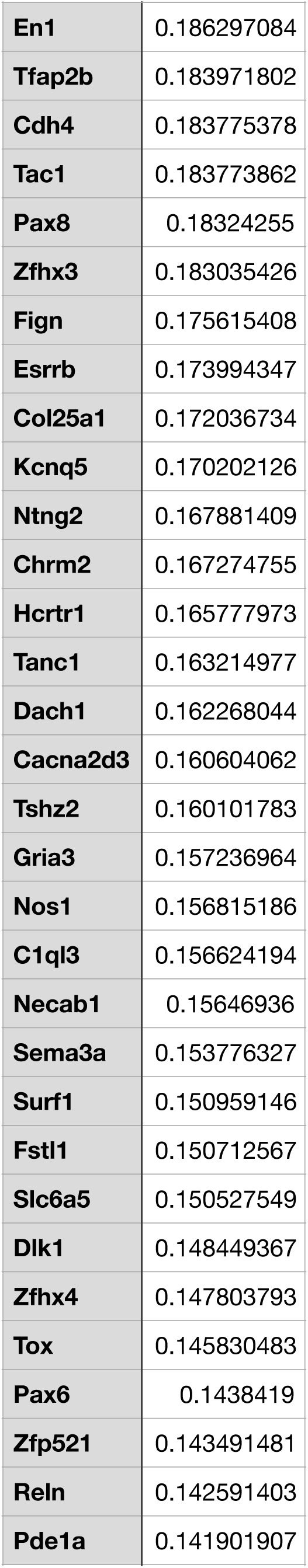
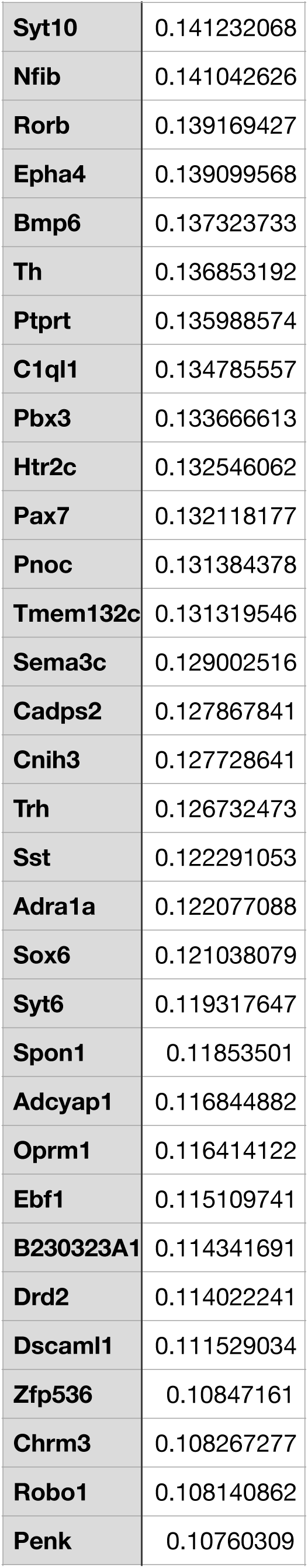
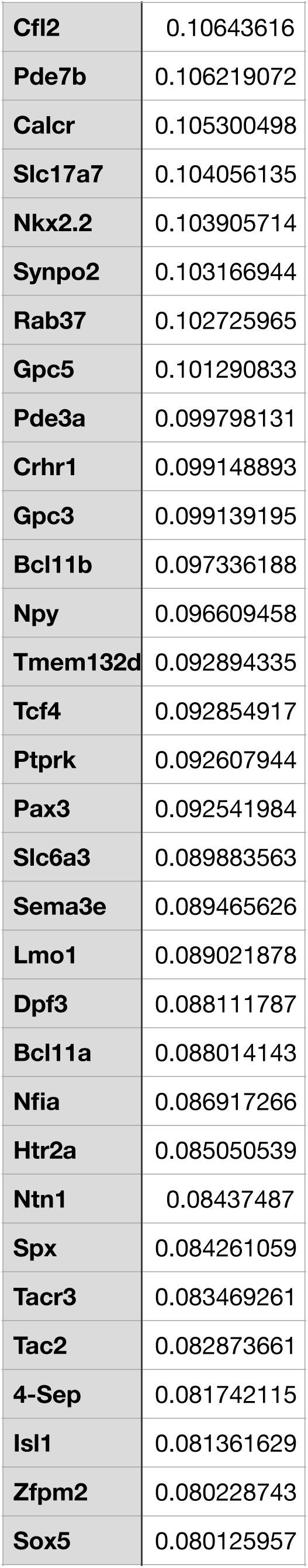
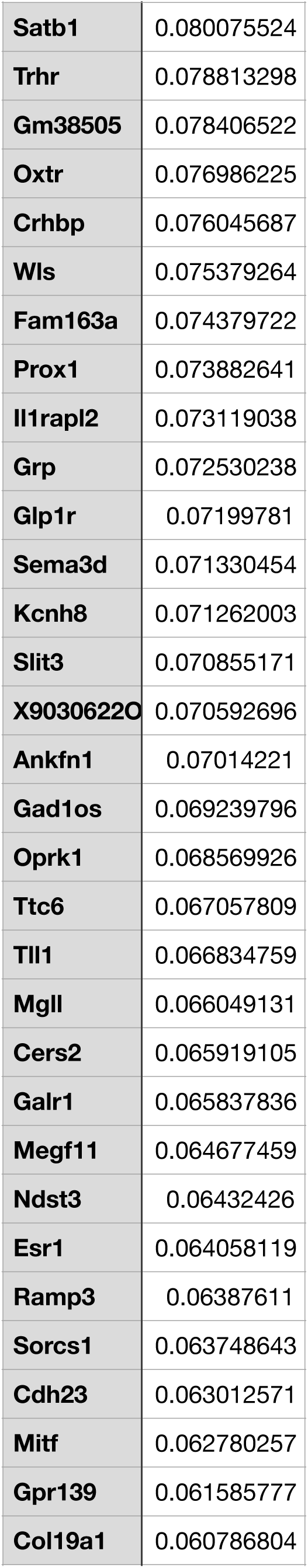
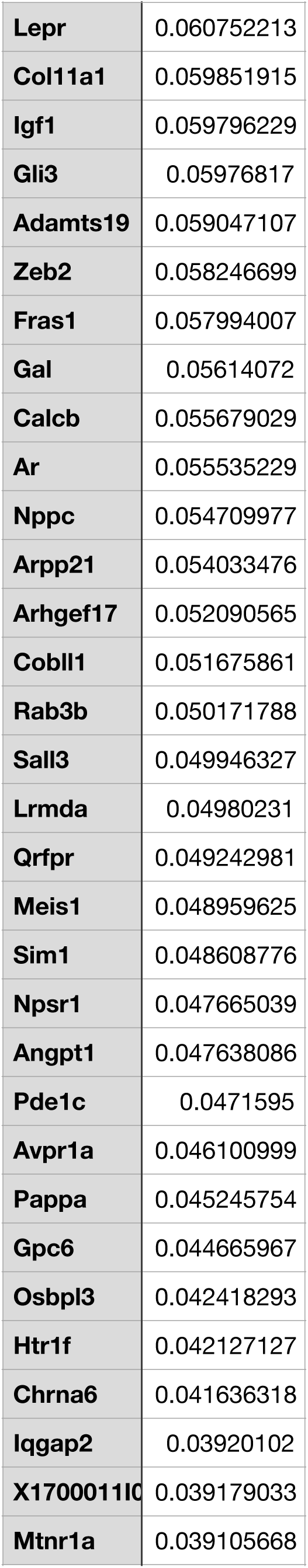
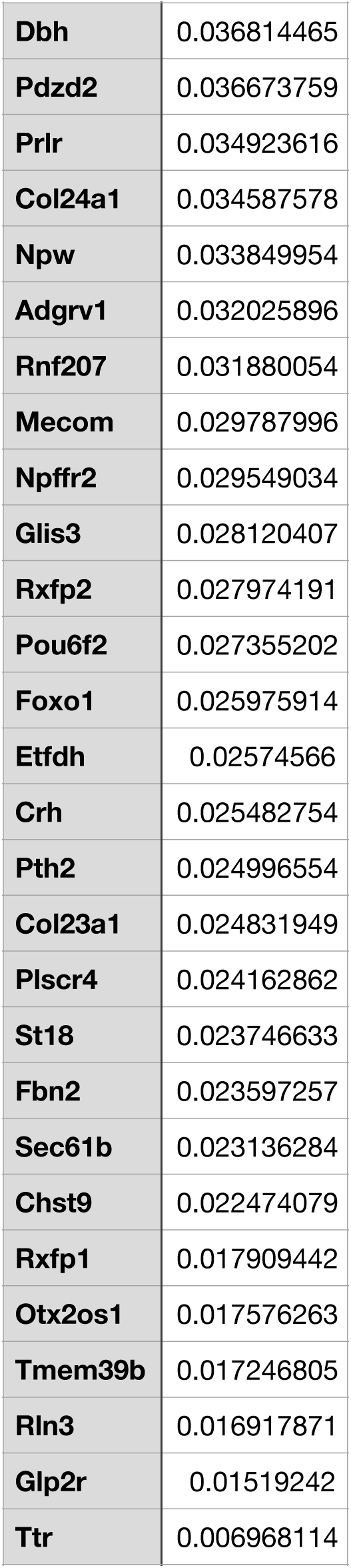

### PAG involvement in behavior and autonomic functions

Spatially restricted lesions and recordings as well as electrical and pharmacological manipulations have attributed distinct behavioral and physiological functions to specific columnar coordinates of the PAG. However, these studies typically did not reach cell type resolution (Deng et al., 2016; Rossier et al., 2021; Tovote et al., 2016). Building upon the vast cell type heterogeneity and complex spatial distribution uncovered by our study, we sought to determine how distinct PAG neuronal populations are involved in various instinctive behaviors. For this purpose, we monitored the expression of multiple IEGs after a variety of behavioral episodes: exposure to a novel object (control), parenting in mothers, fathers, and virgin females, adult- and infant-directed aggression in males, mating, voluntary exercise, and predator exposure (fear and defensive behavior) (Fig 4A; Methods). From an initial panel of 26 IEGs, we further selected seven that displayed strong, correlated, behavior-dependent signal: Fos, Egr1, Fosl2, Fosb, Jun, Nr4a1, and Nr4a3. Monitoring multiple IEGs at single-molecule resolution (IEG Score) provided a high level of robustness and sensitivity. Likely due to highly variable baseline firing rates in the PAG (Wright and McDannald, 2019), many clusters displayed high baseline IEG Scores in controls and across all behaviors (Fig. 4B), prompting us to assess differences in IEG Scores across behaviors within each cluster. Hence, we measured the weighted fold change (WTFC), defined as the relative fold change of the cluster’s mean IEG Score for a given behavior compared to the mean IEG Score for all behaviors in that cluster (Methods).

**Figure 4:**
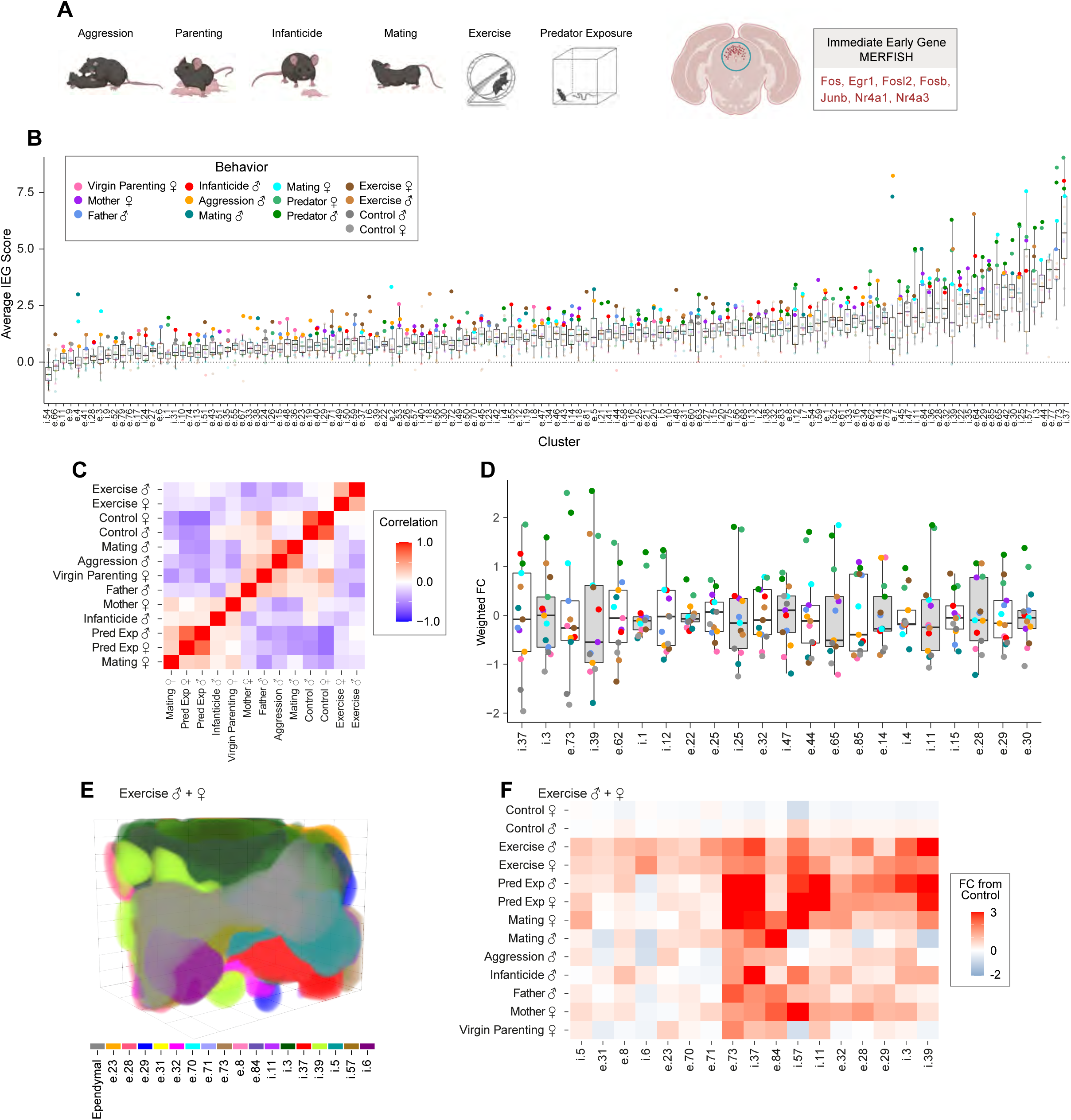
Activation of Neurons during Instinctive Behaviors. A: Overview of behaviors assayed. B: Box plots of mean IEG score for each cluster (columns) for each behavior (colored). Sorted left to right by average IEG score. Dashed line at IEG Score = 0. C: Pearson correlation heatmap of behavior calculated using the weighted fold-change for each cluster and behavior. D: Weighted fold change of select clusters activated by predator exposure, colored as in C. E: 3D organization of clusters activated by exercise. F: Heatmap of weighted fold changes for clusters activated by exercise.

To broadly define the PAG’s involvement across all behavioral responses, we calculated their Pearson correlation using each behavior’s weighted fold change (Fig. 4C). Activity patterns across sexes were closely correlated for controls, exercise, and predator exposure. By contrast, significant differences were found in PAG activity during male and female mating: male mating was mostly correlated to male aggression whereas female mating appeared most correlated with predator exposure. In addition, we found mothers, fathers, and infanticidal males, but not virgin female parenting, to be moderately correlated with predator exposure, suggesting unexpected commonalities between these behaviors.

In addition to identifying the cell clusters that were activated during individual behaviors, we also determined the SMCs that were activated during these behaviors. Notably, individual behaviors preferentially activated only a small subset of SMCs (Fig. S4E, F), by contrast to the SMCs determined from the randomization control described above that were not preferentially activated by any specific behavior. These results support the notion that SMCs represent functionally distinct groups underlying the PAG’s role in various instinctive behaviors.

The PAG stands as a nexus between brain structures involved in sensory processing and internal representation such as the cortex, hypothalamus, or amygdala, and effector circuits within the brainstem. Accordingly, neuronal populations within PAG may encode a specific behavior output or discrete functional elements common to multiple behaviors (i.e. running as part of escaping, pursuing a mate, or a prey). For example, i.1, a dlPAG neuronal population marked by Pax7 and Tfap2b, showed very specific activation after predator exposure (Fig. 4 D), a finding corroborating the involvement of dlPAG in threat assessment (Deng et al., 2016; Esteban Masferrer et al., 2020). By contrast, e.85, representing urocortinergic cells of the Edinger Westphal nucleus, was significantly activated by predator exposure, mothering, fathering, female exercise, and female mating. Overall, male and female predator exposure triggered significant activations in 23 clusters (17.4% of neurons), while male and female parenting activated 11 (6.7% of neurons); male and female exercise, 6 (7.3% of neurons); and male and female mating, 5 (3.5% neurons), reflecting a particularly high engagement of the PAG in defensive behavior and fear responses.

In addition to its central function in directing instinctive behaviors, the PAG also plays a critical role in regulating autonomic output such as breathing rate (Subramanian et al., 2008), heart rate (Lopez-Gonzalez et al., 2020), and blood pressure (Carrive, 1993). In an effort to uncover cell types associated with these functions, we assayed mice after an hour of habituated voluntary exercise on a running wheel. Of the 144 identified neuronal clusters, 17 clusters from several regions of the PAG showed statistically significant increases in IEG score after exercise in males and females relative to controls (Fig. 4 E and F). We interpret their recruitment as an involvement in autonomic and motor functions, though exceptions may exist. Accordingly, many clusters activated by exercise expressed high levels of the metabolism-related receptors Trhr, Adra1a, and Glp1r, and were located dorsally in regions known to cause increases in heart and breathing rate (Lopez-Gonzalez et al., 2020; Subramanian et al., 2008). Consistent with common increases in sympathetic activity, many exercise-activated clusters were also activated by predator exposure. For example, e.28 and e.29 were activated during exercise and predator exposure (Fig. 4D and E) and were enriched for expression of glucagon-like peptide 1 (Glp1) (Ghosal et al., 2013) and Trh (Joseph-Bravo et al., 2015) receptors, supporting a role in the physiological stress response. Intriguingly, these clusters were also activated in mothers and fathers, albeit to a lesser degree. As e.29 is the only dorsal cluster expressing Oxtr and e.28 expresses Galr1, these clusters represent potential targets of galanin (Kohl et al., 2018; Wu et al., 2014) and oxytocin (Schiavo et al., 2020; Scott et al., 2015) – expressing neurons shown to be critical to parental behavior. Interestingly, administration of oxytocin into dorsal PAG induces anxiolytic effects (De Oliveira Sergio et al., 2020), suggesting a circuit mechanism linking the modulation of the stress response and parenting.

### Discriminating function from spatially intermingled dorsal and lateral PAG cells

The dorsal and lateral regions of the PAG have been proposed to mediate active coping strategies displayed during fear and aggression, yet, whether these actions are carried out by the same cells, and whether other behavior functions are performed in these regions remains unknown. Consistent with the literature, we observed that predator exposure and aggression preferentially activated clusters in dorsal and lateral PAG (Bandler and Keay, 1991; Evans et al., 2018; Falkner et al., 2020) (Fig. 5A, B), with predator exposure additionally activating several more ventral clusters(Tovote et al., 2016) (Fig. 5B). However, unexpectedly, we also found virgin female parenting activated seventeen clusters solely in dorsal PAG. Fourteen of these 17 clusters were commonly activated by aggression but only one by predator exposure (Fig. 5 C, D). As the virgin females assayed only retrieved and never attacked pups, such conservation with aggression likely indicates a non-motor role for these populations. In contrast to the common activation of clusters during aggression and virgin female parenting, and despite considerable spatial overlap, only three clusters were activated by both aggression and predator exposure (Fig. 5 D-G). While both predator exposure and aggression recruited clusters from dorsal SMCs 2, 4, and 6, aggression specifically recruited SMCs 7, 9, 10, 13, and 14; virgin female parenting preferentially activating clusters from SMCs 2, 9, and 13 (Fig. S4 E and G).

**Figure 5:**
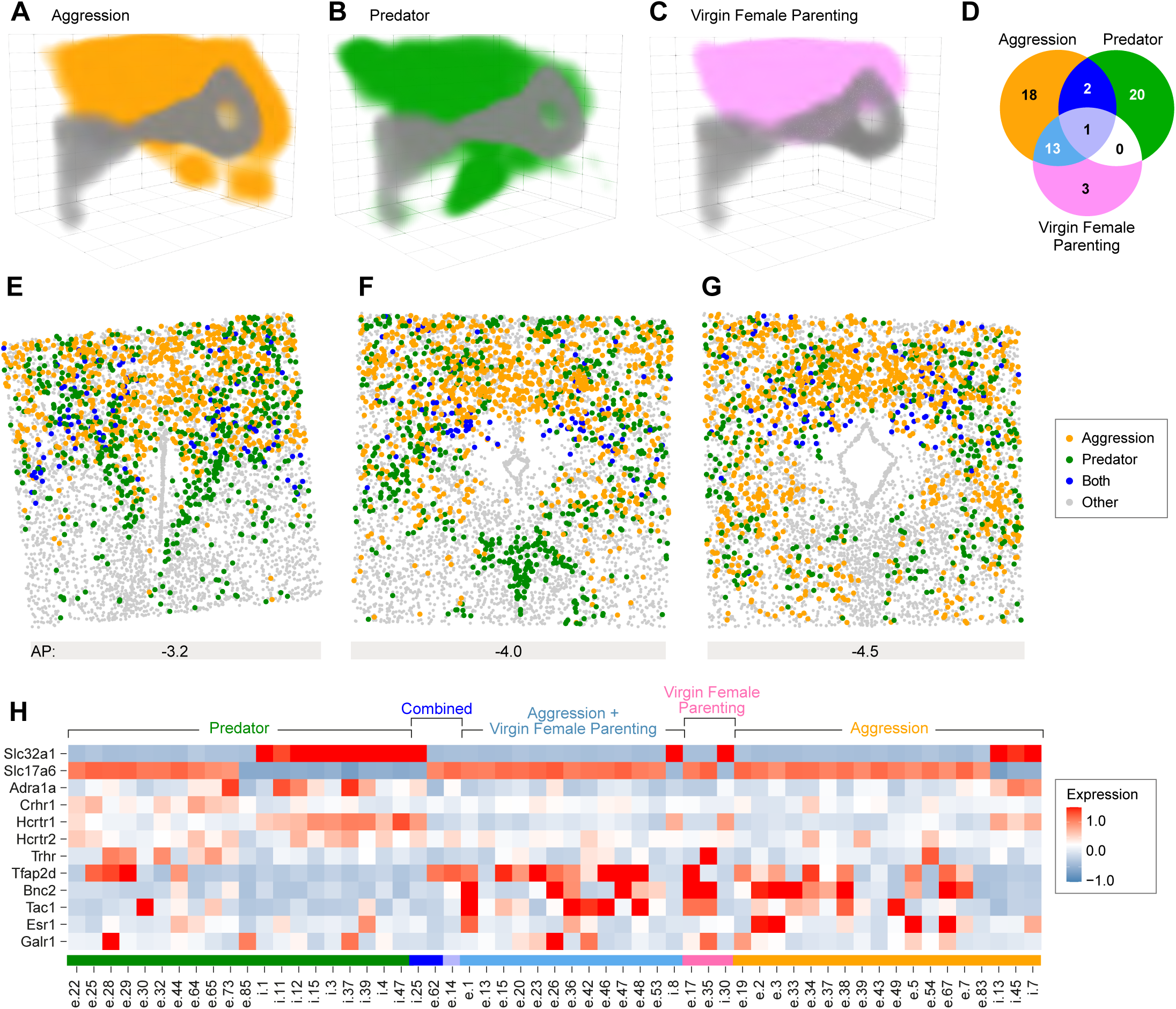
Distinct Behavior Profiles from Overlapping Neuronal Populations. A-C: 3D distribution of clusters activated by aggression, fear, and virgin female parenting, respectively. D: Venn diagram of clusters specifically activated by aggression, predator exposure, or virgin female parenting, and their overlap. E-G: Coronal sections from different anterior-posterior locations showing intermingled clusters activated by aggression, predator exposure, or both. H: Z-score expression heatmap of predator exposure, virgin female parenting, or aggression activated clusters.

Although populations activated by predator exposure, aggression, and virgin female parenting were largely intermingled, specific molecular features segregated them. Clusters activated by aggression and virgin female parenting were mostly excitatory, often enriched for expression of Tac1, Galr1, and the transcription factors Tfap2d and Bnc2, while several aggression-only clusters expressed Esr1 (Fig. 5H). Poised to modulate fear or anxiety, we found enrichment of the orexin receptor genes Hcrtr1 and Hcrtr2, the corticotropin releasing hormone receptor Crhr1, and the alpha-adrenergic receptor gene Adra1a in clusters activated by predator exposure (Fig. 5H). Consistent with this notion, activation of Hcrtr1 (Pourrahimi et al., 2019) and Crhr1 (Sergio Tde et al., 2014) expressing clusters in PAG has been reported to induce both heightened anxiety and antinociception. Additionally, Hcrtr1/2 (Annerbrink et al., 2011; Gottschalk et al., 2019), Crhr1 (Weber et al., 2016), and Adra1a (Zhang et al., 2017) polymorphisms are strongly associated with human panic disorders and targets for clinical therapy (Johnson et al., 2010; Lohoff et al., 2013), suggesting cells expressing these genes may be of critical importance to the PAG’s role in anxiety (Fanselow, 1991). Thus, these results molecularly delineate spatially overlapping, yet mostly mutually exclusive cell populations critical to predator exposure and aggressive or virgin female parenting behavior.

### Cell types involved in parenting or infanticide

In mice, mothers and fathers robustly display parental displays such as retrieval and licking, while virgin females usually retrieve pups, but with higher latencies and less robust parenting (Kohl and Dulac, 2018). By contrast, virgin males often show lethal aggression towards pups (infanticide) (Kohl and Dulac, 2018). The PAG has previously been implicated in regulating aspects of parenting (Kohl et al., 2018; Lonstein and Stern, 1997a, b; Stern and Lonstein, 2001; Sukikara et al., 2006; Zhang et al., 2021). It receives dense input from the medial preoptic area (POA) and optogenetically stimulating galanin expressing POA (POAGal) terminals in rostral PAG (rostral PAG) increases pup grooming(Kohl et al., 2018). Hence, we sought to define PAG cell types involved in infant-evoked behavior. In response to pups, we observed several types of cluster activities: significant activity during mother and/or father parenting (Fig. 6A; light blue), active in virgin female parenting (pink), in infanticidal males (red), or, to our surprise, commonly active in mothers and infanticidal males (purple; discussed below).

**Figure 6:**
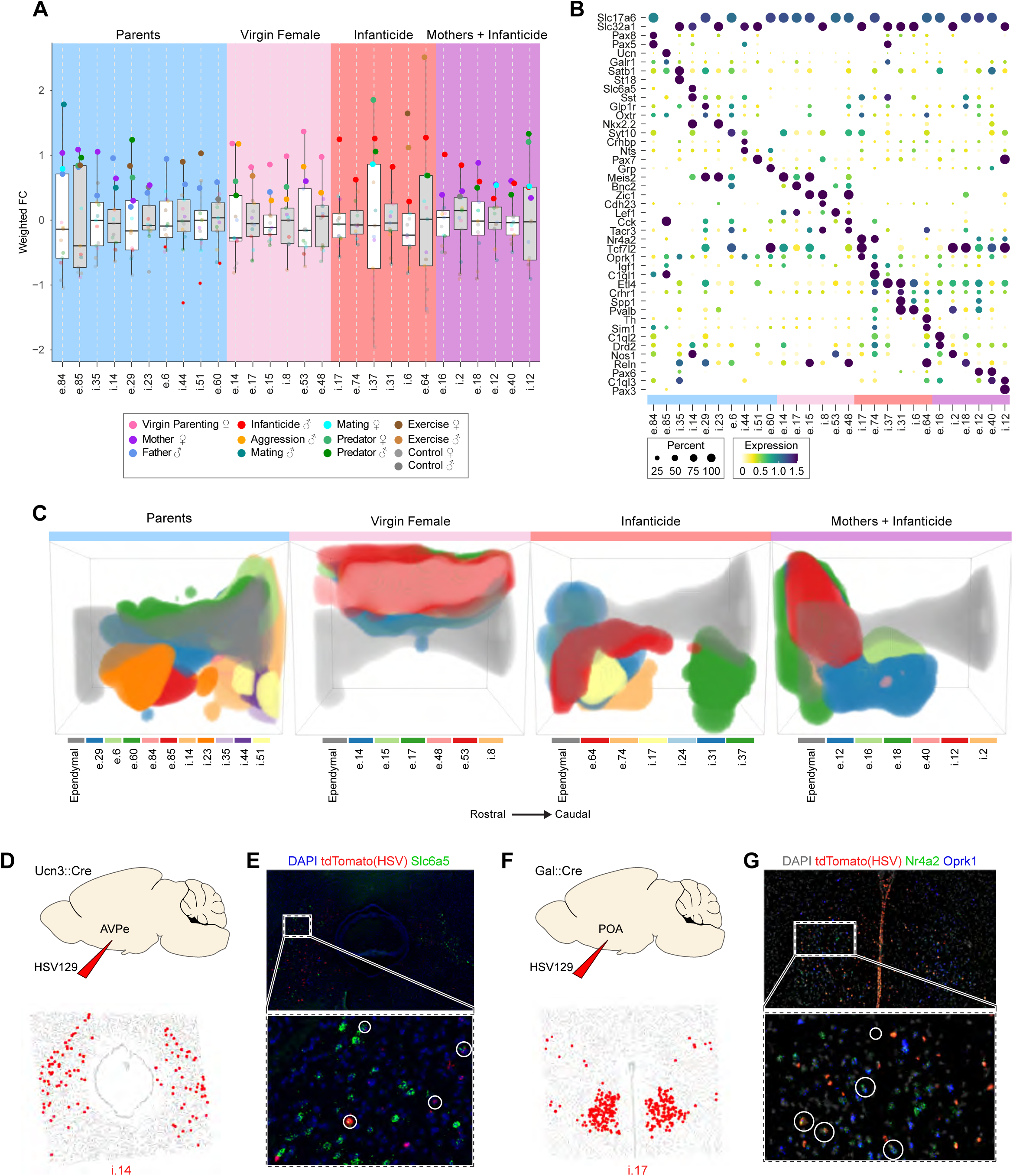
Regional Recruitment of PAG Neurons by Pup-related Behaviors. A: Box plot of parenting or infanticide related clusters. Light blue: mother and father active, right four father active; pink: virgin female active; red: infanticide active; purple: mother and infanticide active. For emphasis, significantly activated clusters are plotted at full opacity while others are shrunken at reduced opacity. B: Expression matrix of clusters in A where dot size represents percent of cells expressing gene and color represents mean cluster z-score. C: Sagittal three-dimensional distribution of clusters for each group detailed in A. D: Diagram of HSV129 injection into AvPe of Ucn3::Cre mice (top) and distribution of parenting-active i.14 in caudal PAG (bottom). E: Example RNAScope in situ from of anterogradely tdTomato labeled HSV cells in caudal PAG. Yellow circles detail Slc6a5 and HSV co-labeled cells. F-G: Same as in D and E but using Gal::Cre mice targeting i.17 with Nr4a2 and Oprk1 in rostral PAG.

We selected the top six most selectively activated clusters for each condition. Many parenting clusters were marked by the transcription factor Satb1 and expressed moderate to significant amounts of Galr1, Oxtr, and Oprk1, indicating that galanin (Kohl et al., 2018; Moffitt et al., 2018; Wu et al., 2014), oxytocin (Schiavo et al., 2020) and opioids (Mann et al., 1991; Sukikara et al., 2006) may exhibit their effects on parenting partially via these cells (Fig. 6B). By comparison, virgin female parenting clusters expressed scant amounts of Galr1, Oxtr, and were instead marked by Meis2, Bnc2, and Zic1 (Fig. 6B).

Remarkably, when we expanded our selection to include four clusters uniquely activated in fathers (e.6, i.44, i.51, e.60), we noted depleted activity, or potentially inhibition, by infanticide in these clusters (IEG Scores substantially below that of controls and other behaviors), suggesting that infanticidal behavior may act in part by suppressing cell types activated during paternal behavior (Fig. 6A). Clusters active during infanticide often expressed Etl4, Pvalb, Crhr1 and Oprk1 (Fig. 6B). As injection of Crhr1- selective antagonist into dorsal PAG is anxiogenic (Litvin et al., 2007), and more broadly, considering Crhr1’s role in stress (Weninger et al., 1999), these populations may reflect the documented activation of stress responses in infanticidal mice (Zhang et al., 2021).

The PAG is thought to integrate diverse information about internal state and environment to select the appropriate adaptive behavior (Sukikara et al., 2006). Selection between mutually incompatible behaviors (ex. flight and parenting) requires the activation and suppression of antagonistic circuits within the PAG, which are often spatially segregated (Sukikara et al., 2006, 2010; Tovote et al., 2016). To decipher the spatial logic of pup-related behaviors in the PAG, we plotted each of the four groups described above (mother/father parenting, virgin female parenting, infanticide, mother parenting + infanticide) in 3D space (Fig. 6 C). Strikingly, we found that populations activated during mother and father parenting, virgin female parenting, and virgin male infanticide occupied largely non-overlapping space. Clusters activated during mother and father parenting were located caudally in regions necessary for kyphosis (Stern and Lonstein, 2001) and receiving input from anterior hypothalamic structures critical to parental behavior. Accordingly, we hypothesized that some of these clusters may receive input from a previously described Brs3- and Unc3-expressing excitatory population of the anteroventral periventricular nucleus (AvPe) (cluster E1) that is only active during parental behavior of mothers (Moffitt et al., 2018). To assess this connectivity, we injected the conditional, anterograde virus H129ΔTK-TT (H129, expressing tdTomato) into the AvPe of Ucn3-Cre mice (Lo and Anderson, 2011) (Fig. 6D). Indeed, 48-60 hours after injection, we found the cluster i.14 activated by parenting to be colabelled with tdTomato and Slc6a5 in caudal PAG (Fig. 6E). As the caudal PAG region is critical for kyphosis (Lonstein and Stern, 1997a) and projects to medullary premotor neurons (Daniels et al., 1999), these data indicate AvPe and caudal PAG may form the previously hypothesized feedforward circuit activating medullary premotor neurons (Lonstein and De Vries, 2000). Surprisingly, virgin female parenting did not activate caudal PAG, but instead dorsal SMCs 13, 2, and 9 (Fig 6C; pink; Fig. S4E and S4G), regions otherwise associated with assessment (Deng et al., 2016; Esteban Masferrer et al., 2020), defense (Deng et al., 2016; Siegel et al., 1997), and fear (Esteban Masferrer et al., 2020; Tovote et al., 2016). Peculiarly, these clusters often showed significant activation by aggression, indicating that while virgin females retrieve pups, the neural processing of this function involves some circuits not active in mothers and fathers.

Of 38 clusters activated during infanticide, 25 were located rostrally in SMCs 5, 18, 4, 11, and 8, while a subset of clusters common to mother parenting and infanticide were preferentially found in SMCs 18, 4, and 15 (n=12/18; Fig. 6C; Fig S4E and S4G). Intriguingly, several clusters common to mother parenting and male infanticide were also significantly activated during predator exposure and located rostrodorsally in a region previously shown to be necessary for mothers to shift from parenting behavior to threat assessment during predator exposure (Sukikara et al., 2010). Accordingly, the activation of the SMC 18 clusters e.18, e.40, and i.12 by mothers, infanticide, and predator exposure suggests a potential role in assessment that supersedes either parenting or infanticide. As we previously demonstrated, inhibitory rostolateral neurons receive input from POAGal neurons involved in parenting (Kohl et al., 2018), prompting us to hypothesize that the rostrolateral PAG clusters i.17 and i.31 activated by infanticide may be the targets of POAGal neurons. Indeed, 48-60 hours after injection, anterograde H129 tracing from POAGal neurons labeled i.17 neurons marked by Nr4a2 and Oprk1 (Fig. 6F, G) as well as i.2 neurons marked by Slc32a1, Tcf7l2, and Nos1 (Fig. S5). This mostly inhibitory connection between parental POAGal neurons (Moffitt et al., 2018) and rostrolateral PAG, a region critical for predation (Sukikara et al., 2006; Tulogdi et al., 2015), puts forth a model whereby POAGal neurons act in part by inhibiting predatory circuits that are activated during infanticide. Indeed, recent findings have demonstrated a substantia innominata-PAG circuit that indiscriminately initiates both predation and infanticide (Zhu et al., 2021). Collectively, these data indicate specific PAG regions play a common role in the initiation of appropriate pup-related responses and further suggest a prominent role for local PAG circuitry in behavior selection (Michael et al., 2020; Tovote et al., 2016; Vaaga et al., 2020).

### Spatial activation of aggression and mating related cell types

We next sought to decipher the neural basis of another set of behaviors: aggression and mating. The PAG is known to be critical for initiating aggression (Falkner et al., 2020; Mos et al., 1982; Siegel et al., 1997) and mating (Hashikawa et al., 2016; Sakuma and Pfaff, 1979; Yamada and Kawata, 2014), however, due in part to its broad activation and wide-ranging functionality, it has remained unclear which specific neuronal populations are involved in each or both of these two behaviors (Hashikawa et al., 2016). Likely representing a common output between behaviors, we observed that many clusters significantly activated by aggression were also active during male and sometimes female mating (Fig. 7A). Of 34 aggression- and 19 male mating-activated clusters, 9 were co-activated by both behaviors, localized to the dorsal and lateral regions (Fig. 7B). Seven of the nine were found in SMCs 14, 10, and 2 (Fig. S4E and S4G). Five caudal PAG clusters, along with the serotonergic dorsal raphe (e.70) showed common activation between male and female mating, consistent with previous literature (Fig. 7C). Predator exposure showed common activity with 8/25 female mating-activated clusters found in rostral and ventral zones (Fig. 7D). More selectively, three other caudal PAG/dorsal raphe clusters were found to be activated by male mating, but not aggression, female mating, or predator exposure: i.19 and e.71, also activated by exercise (Fig. 7A), and i.14, an SST-marked glycinergic population also activated by mothering and fathering (Fig. 6A). In contrast, while female mating significantly activated two clusters (e.12 and i.16) that were not activated by male mating or aggression, these clusters were still mildly activated by predator exposure in males and females. Hence, we discover several molecularly discrete cell types that are recruited by mating, as well as other behaviors that may hint at their specific function. Notably, many of the clusters activated during both aggression and mating (male and female) were Esr1+ neurons (Fig. 7E). Further, the organization of the Esr1+ clusters in a triangular distribution in l- and dmPAG was strikingly similar to projection patterns the PAG receives from Esr1+ cells of the ventrolateral-ventromedial hypothalamus (VMHvl) (Falkner et al., 2020) (Fig. 7F and G).

**Figure 7:**
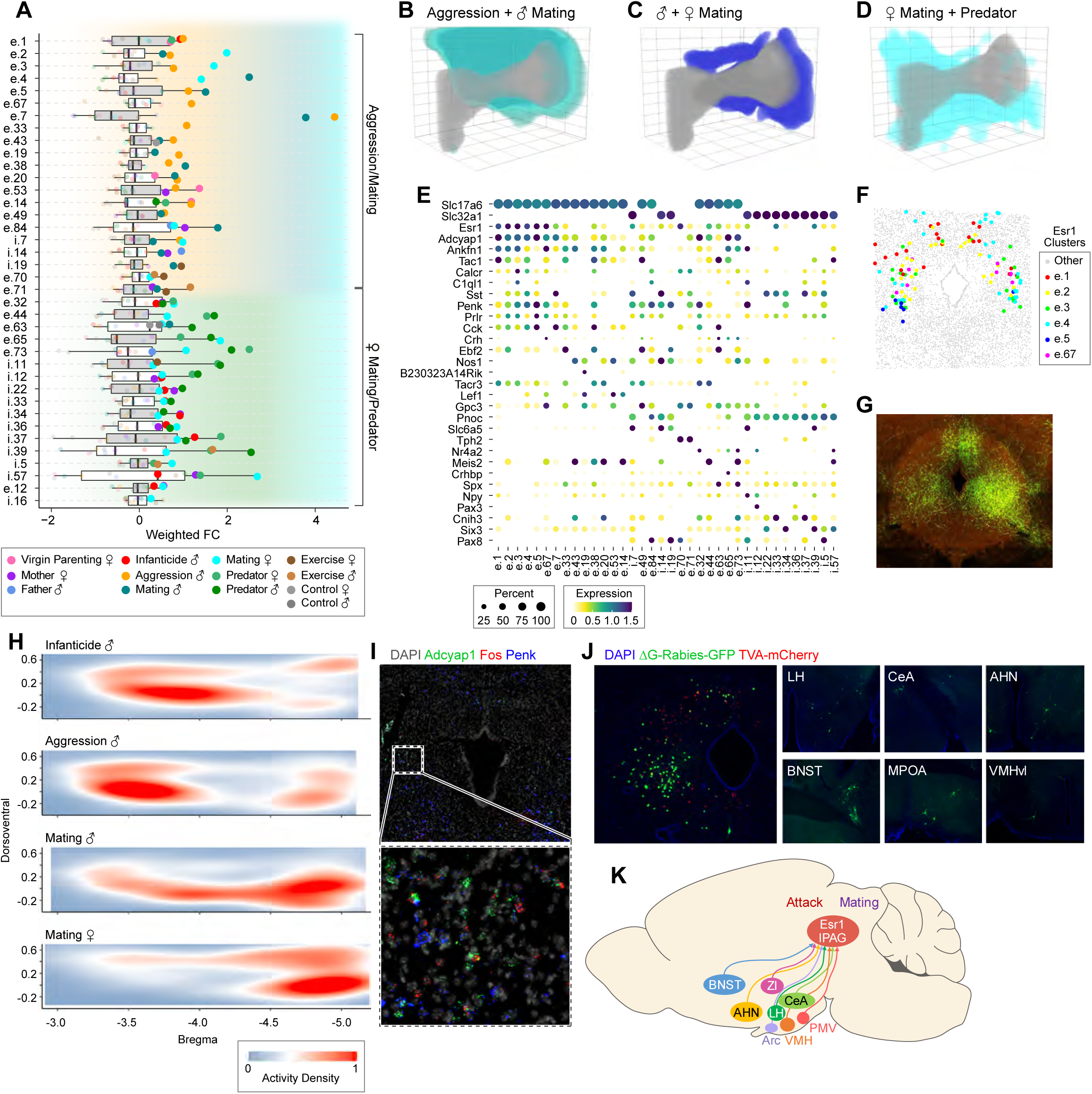
Intra-cluster Differences in Spatial Activation of Esr1 Neurons. A: Boxplot of aggression or mating activated clusters or female mating and fear activated clusters. For emphasis, significantly activated clusters are plotted at full opacity while others are shrunken at reduced opacity. B-D: 3D diagrams depicting the spatial distribution of cells activated by overlapping behaviors. E: Expression matrix of clusters in A where dot size represents percent of cells expressing gene and color represents mean cluster z-score. F: Coronal distribution of Esr1+ clusters. G: Coronal section of the PAG showing conditional labeling from Esr1 cells of the VMHvl, from Allen Brain Institute. H: From top to bottom, sagittal views of Esr1+ cluster activity across space in infanticide, aggression, male mating, and female mating. I: RNAScope in situ of postnatal day 3 pup PAG after induction of vocalization. J: Monosynaptic retrograde tracing using g-deleted rabies-GFP and TVA-mCherry in Esr1::Cre mice. K: Diagram of socially relevant inputs into lPAG Esr1+ PAG cells.

Previous work in rats and cats identified rostral PAG as a potential locus for aggression (Bandler and Carrive, 1988; Siegel et al., 1997), while caudal PAG activation facilitates female mating (Sakuma and Pfaff, 1979). Furthermore, the PAG receives differential inputs along the rostrocaudal axis: the lumbar spinal cord projects densely to caudal PAG while the cervical enlargement projects progressively more to rostral PAG (Bandler and Shipley, 1994). Accordingly, we reasoned that, with many Esr1+ clusters spread across the full ∼2.5mm length of the PAG, their activation along this axis may vary. Overall, male/female mating and aggression activated all regions that include Esr1+ clusters more than other behaviors. However, within individual Esr1+ clusters, we found that aggression preferentially activated the rostral half, while conversely, female mating activated the caudal half of the same cluster (Fig. 7H; Fig. S6A). Remarkably, male mating recruited cells common to both aggression and female mating, displaying intermediate levels of activation in both the rostral and caudal cells. Like aggression, infanticide also preferentially activated the rostral cells of e.1 (Fig. 7 A and H; Fig. S6A), which remained unchanged in parents (relative to controls). We initially observed these spatial differences within one large excitatory Esr1+ cluster and suspected the differences may be due to a lack of thorough clustering, and as such, resolved this large cluster out into seven stable clusters. While e.5 was found preferentially in caudal PAG, the remaining six were still found spread across the length of the PAG, suggesting that observed spatial activation differences are indeed taking place within molecularly defined cell types. This difference in spatial activation suggests that rostral PAG Esr1+ cells are involved in the manifestation of aggressive or pursuit behaviors, whereas caudal PAG Esr1+ cells are involved in mating behavior. Moreover, this difference emphasizes the substantial impact played by cell location in function, even within a molecularly well-defined cell population.

Among Esr1+ clusters, e.4 had extremely specific activity in male mating (Fig. 7A). Additionally, while its IEG scores for other behaviors hovered around zero, male and female predator exposure both had a negative IEG Score for this cluster. Male mice emit PAG-dependent ultrasonic vocalizations in the presence of a female (Tschida et al., 2019), while inhibitory amygdala to PAG projections have been shown to directly suppress vocalizations, presumably critical for contexts when silence is required, such as during predator exposure (Michael et al., 2020). We reasoned that these Esr1+ e.4 cells could represent neurons responsible for vocalization based on its activation during male mating, slight activation during female mating (females often vocalize in response to male pursuit), position in the PAG (Chen et al., 2021; Tschida et al., 2019), and potential inhibition during predator exposure. We tested this hypothesis in a non-social setting by isolating postnatal day 3 pups from their mother and robustly inducing vocalizations (Fig. S6B). Labeling with Fos and e.4 markers Adcyap1 and Penk showed strong overlap for very specific caudal lPAG cells (Fig 7I). Taken together, these experiments suggest e.4 represents a cell type involved in vocalization from birth onward and that these cells may be inhibited contextually by predator exposure.

Lastly, given the activity defined, we sought to place Esr1+ clusters within a broader context by assessing their neural inputs. lPAG is known to receive input from VMH neurons involved in mating and aggression (Falkner et al., 2020), lateral hypothalamus (LH) (Li et al., 2018) and CamkIIa-POA neurons involved in pursuit (Park et al., 2018), as well as POA and CeA inputs critical for vocalization (Chen et al., 2021; Michael et al., 2020). Hence, to assess their connectivity with the above, we performed monosynaptic retrograde rabies tracing (Wickersham et al., 2007) from lPAG of Esr1-Cre mice (n=3).

Consistent with the above notion, GFP+ rabies labeling was detected in several socially relevant brain structures, including anterolateral bed nucleus of the stria terminalis, LH, POA, CeA, VMHvl, and the anterior hypothalamic nucleus (Fig 7 J and K). We further confirmed Esr1+ VMHvl cells project to lPAG Esr1+ cells by performing conditional anterograde H129 tracing and RNAScope validation from the former in Esr1-Cre mice (Fig. S6C). These results identify Esr1+ l/dmPAG neurons as a hub for socially relevant circuits critical to mating and aggression.

## Discussion

Here, we explored a diverse panoply of instinctive functions in a previously poorly parcellated structure, the mouse periaqueductal gray matter. Using snRNA-seq and spatially resolved single-cell transcriptomic measurements by MERFISH we mapped transcriptional and behavioral features in individual cells of the PAG. The high sensitivity of MERFISH (Moffitt et al., 2018) enabled our analysis to capture a wide array of specific neuromodulatory gene expression patterns across the 144 clusters defined.

We identified neuromodulatory genes that were preferentially expressed in excitatory or inhibitory neurons, while many others showed more widespread expression (Fig S1E). Clusters defined by expression of signaling molecules such as Tac1, Adcyap1, Penk, Crh, Pnoc, Spx, Sst, Cck, Nts, Trh, Npy and receptors like Esr1, Prlr, Galr1, Drd2, Tacr3, Trhr were specified throughout the PAG (Figure 8). With these data, a highly granular molecular and cellular parcellation of the PAG emerges that refines and redefines the previous coarse anatomical subdivisions in 4 broad radial columns.

**Figure 8:**
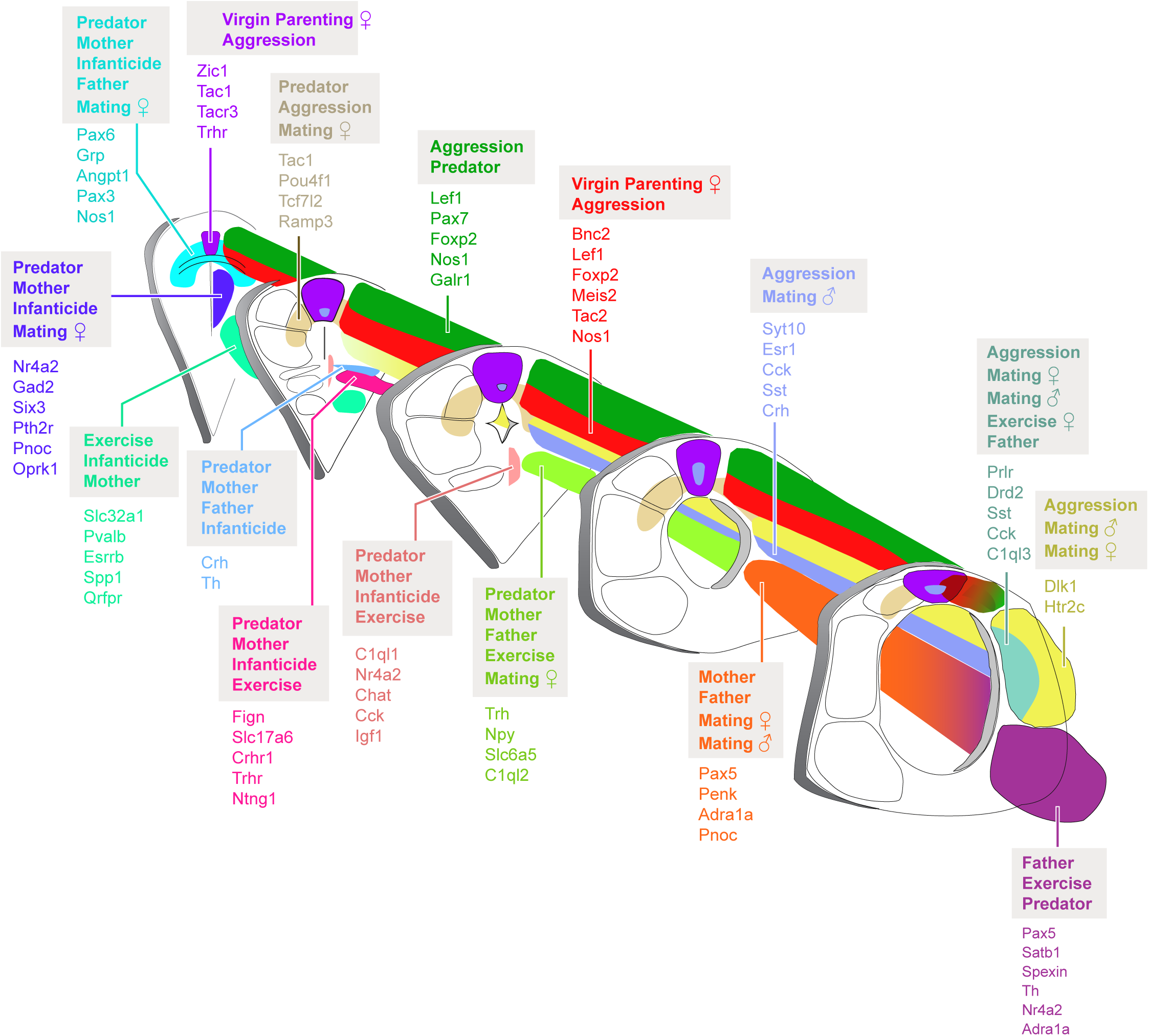
Molecularly Refined Spatial Model of the PAG During Instinctive Behavior. Updated model of PAG distribution using spatial metaclusters as a framework. Colors represent SMCs (excluding non-PAG SMCs). Colored text represents genes highly expressed within corresponding SMCs in diagram. Text in bold represents behaviors that activate a given SMC.

We explored here the precise molecular and anatomical organization of clusters in 3D, providing a comprehensive anatomical framework for transcriptionally defined cell populations in the PAG. These delineations provide substantial insights that may guide the design and interpretation of functional manipulations and recordings along the PAG axes. At a higher level of organization, we define spatial metaclusters (SMCs) that comprise transcriptionally-diverse populations that unite in their spatial distributions. We identify SMCs that agree with classical columnar structures in the PAG (e.g.dmPAG- specific SMC 9) and hence defining the cellular composition of these structures, other that refine the existing columnar structures (e.g. SMCs 2 and 13 representing outer and inner dlPAG groups), and finally some identifying spatial motifs that do not agree with classical columnar structures (e.g. the dorsal-arch SMC 6) (Figure 8). Thus, these newly defined spatial compartments both complement and revise the existing anatomical delineations of the PAG.

We provide spatially and functionally separable families of cells by defining activity of SMCs in response to a common set of or specific behaviors (Figure 8 and S4). For instance, in SMCs 4, 18, and 15, IEG score decreases in exercise, but increases during predator exposure, mothering, and infanticide, substantiating the behavioral induction of and rostral PAG’s role in anxiety (Duman et al., 2008; Singewald and Sharp, 2000; Zhang et al., 2021). At the same time, these data highlight the internal functional diversity of SMCs: aggression and predator exposure both activating four clusters within SMC 6, with only one coactivated. Thus, our molecular parcellation provides critical cell type-specific entry points to investigate their specific function in instinctive responses.

At the single-cluster level, we find a range of behavioral involvements: from being highly specific to one behavior, to being commonly recruited across a set of behaviors. This suggests that at the level of the PAG, information may condensate to specific motor or autonomic actions shared across distinct behaviors. An example of such would be the lPAG-evoked action of pursuit, which is common to offensive acts of aggression (Falkner et al., 2020), male mating (Falkner et al., 2020), and predation (Li et al., 2018; Park et al., 2018). Furthermore, predator exposure and parenting shared heightened cluster activations in e.29 and the Edinger Westphal (EW) urocortinergic e.85, representing potential stress or anxiety-related pathways common in both conditions. Indeed, the principally projecting EW has recently been shown to increase measures of anxiety and be activated by several forms of stress (Priest et al., 2021). Similarly, the coactivation of mating and parenting in caudal vlPAG e.84 cells, which are somatopically connected to lumbrosacral segments of the spinal cord (Bandler and Shipley, 1994), represent their potential involvement in regulating lumbar/hip posture for important caudal PAG related actions like lordosis (Sakuma and Pfaff, 1979), mounting (Holstege, 2016), and kyphosis (Stern and Lonstein, 2001). Lastly, the approach-avoidance model of female parenting proposes that females exhibit parental behavior through hormonal changes or repeated exposure to pups (Stolzenberg and Mayer, 2019). Virgin females of other rodent species often actively avoid pups (Stolzenberg and Mayer, 2019), while lesions to POA in mice increase infanticidal behavior in virgin females or mothers (Tsuneoka et al., 2013). This suggests the presence of pup-aversive circuits that are at least partially engaged in virgin females and could explain the initial hesitance virgin females display upon exposure to pups. Such aversion is corroborated by dorsal clusters coactivated by male aggression and virgin female parenting. Indeed, for e.14, activated by male aggression and virgin female parenting, the additional coactivation of predator exposure in both males and females bolsters the assumption that these cells encode information critical to evaluating potential threat (Reis et al., 2021).

Significantly, we provide evidence that within individual clusters, activity across behaviors may vary substantially along its rostrocaudal axis. For example, we observed that aggression preferentially activated Esr1+ neurons in the rostral part of PAG while female mating preferentially activated the caudal aspect. Interestingly, inhibition of rostral PAG Esr1+ neurons have been reported to increase lordosis (Ishii et al., 2017), suggesting that inhibition of aggression-activated rostral PAG Esr1+ neurons may act to facilitate mating. Hence, our work provides important spatial characteristics of function in the PAG that can be used to more precisely target populations of interest.

Broadly, this work provides a novel framework for discovering three-dimensional spatial motifs in neuronal cell types, while simultaneously defining their functional output. Specifically, we used this method to unmask the transcriptional, spatial, and functional logic of a highly conserved, but poorly understood nexus paramount to instinctive behavior and homeostatic function, the PAG. These data provide a transformative framework for future anatomical, molecular and functional explorations of the PAG.

## Acknowledgments

We thank Dhananjay Bambah-Mukku for guidance during single-cell and tissue preparations, MCB Graphics for help with illustrations and the Harvard Bauer Core Facility for assistance with sequencing and cell sorting. The work was supported by grants R01MH113094 and U19MH114821 from the National Institutes of Health to X.Z and C.D. E.V was supported by the Ruth L. Kirschstein National Research Service Award (NRSA) Individual Predoctoral Fellowship (F31). C.D., and X.Z. are Howard Hughes Medical Institute Investigators.

## Author contributions

E.V., C.D, S.W.E., and X.Z conceived of the study. Experiments and data analyses were performed by E.V and S.W.E with help from W.J. E.V., C.D., and X.Z. wrote the manuscript. Competing interests: X.Z. is an inventor on patents applied for by Harvard University related to MERFISH, and a co-founder and consultant of Vizgen. Data and materials availability: snRNA-seq data are available at the Neuroscience Multiomic Data Archive (NeMO) (ID: dat-diw5xr1). MERFISH data are available at the Brain Image Library (ID: 0a4681ec1919cda5), and the analysis software is available at https://github.com/ZhuangLab/MERFISH_analysis.

## Supplemental Figures

**Figure S1:**
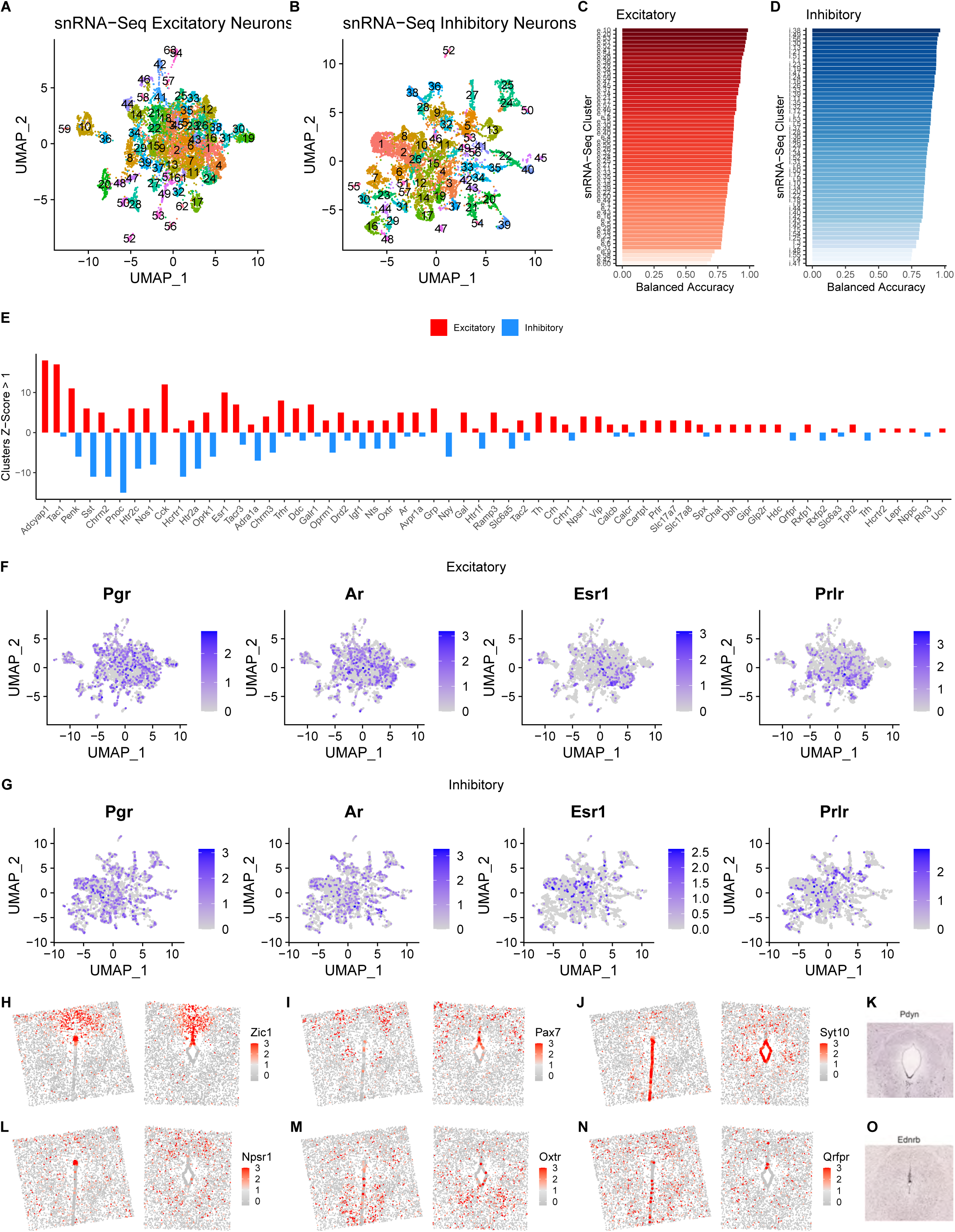
Defining MERFISH Gene Library using snRNA-Seq Clustering, Related to Figure 1 and Table 1. (A and B): Uniform Manifold Approximation Projection (UMAP) display of snRNA-Seq excitatory (A) and inhibitory (B) neuronal clusters. (C and D): Balanced Accuracy per excitatory (C) and inhibitory (D) cluster for SVM predictions made using reduced MERFISH library gene set. (E): Barplot depicting the number of clusters with z-score>1 for each neuromodulatory gene; Up/red indicates number of excitatory clusters and down/blue represent number of inhibitory clusters. (F and G) Log normalized expression of sex hormone receptors in excitatory and inhibitory snRNA-Seq subsets. (H-N): Z-score expression of signaling related genes in ependymal cells using MERFISH. (K and O): Coronal in situ hybridization images of additional genes identified within snRNA-Seq but not included in MERFISH (Images from Allen Brain Atlas).

**Figure S2:**
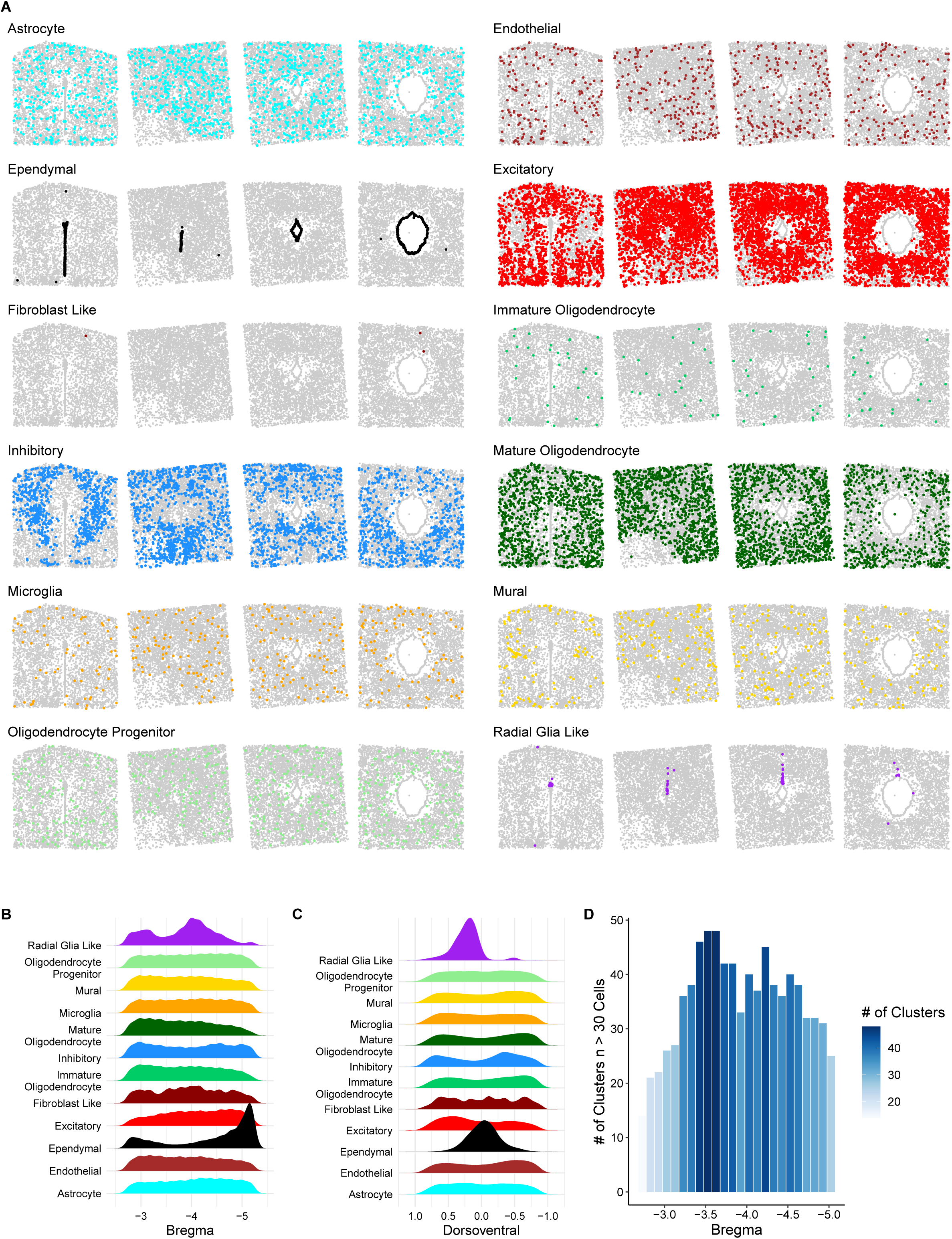
Spatial Distribution of Cell Classes in MERFISH, Related to Figure 2. (A and B): Density distribution each cell class along the rostrocaudal axis (A; read as AP distance from Bregma) and dorsoventral axis (B; read as distance from the per-slice median aqueduct coordinates). (C) Coronal slices displaying inhibitory neuron distribution in rostral (top) and caudal (bottom) regions of the PAG. (D) Distribution of ependymal and radial glia-like cells from rostral to caudal coronal sections. (E): Barplot displaying the number of abundantly represented clusters (n>30) across the rostrocaudal axis.

**Figure S3:**
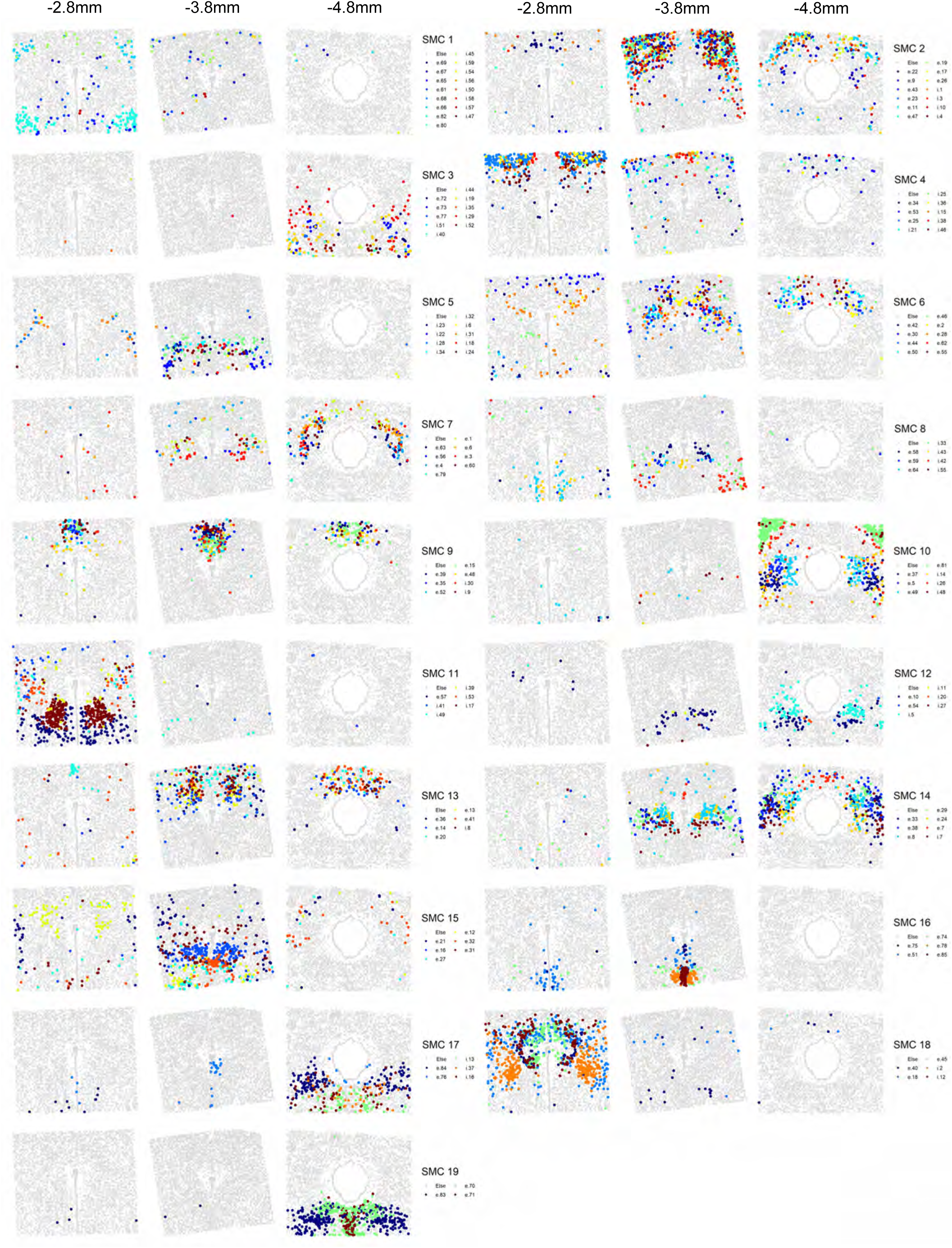
Position of Spatial Metaclusters, Related to Figure 3 and Table 2. Rostral, intermediate, and caudal positions of each SMC where each color represents a cluster within an SMC.

**Figure S4:**
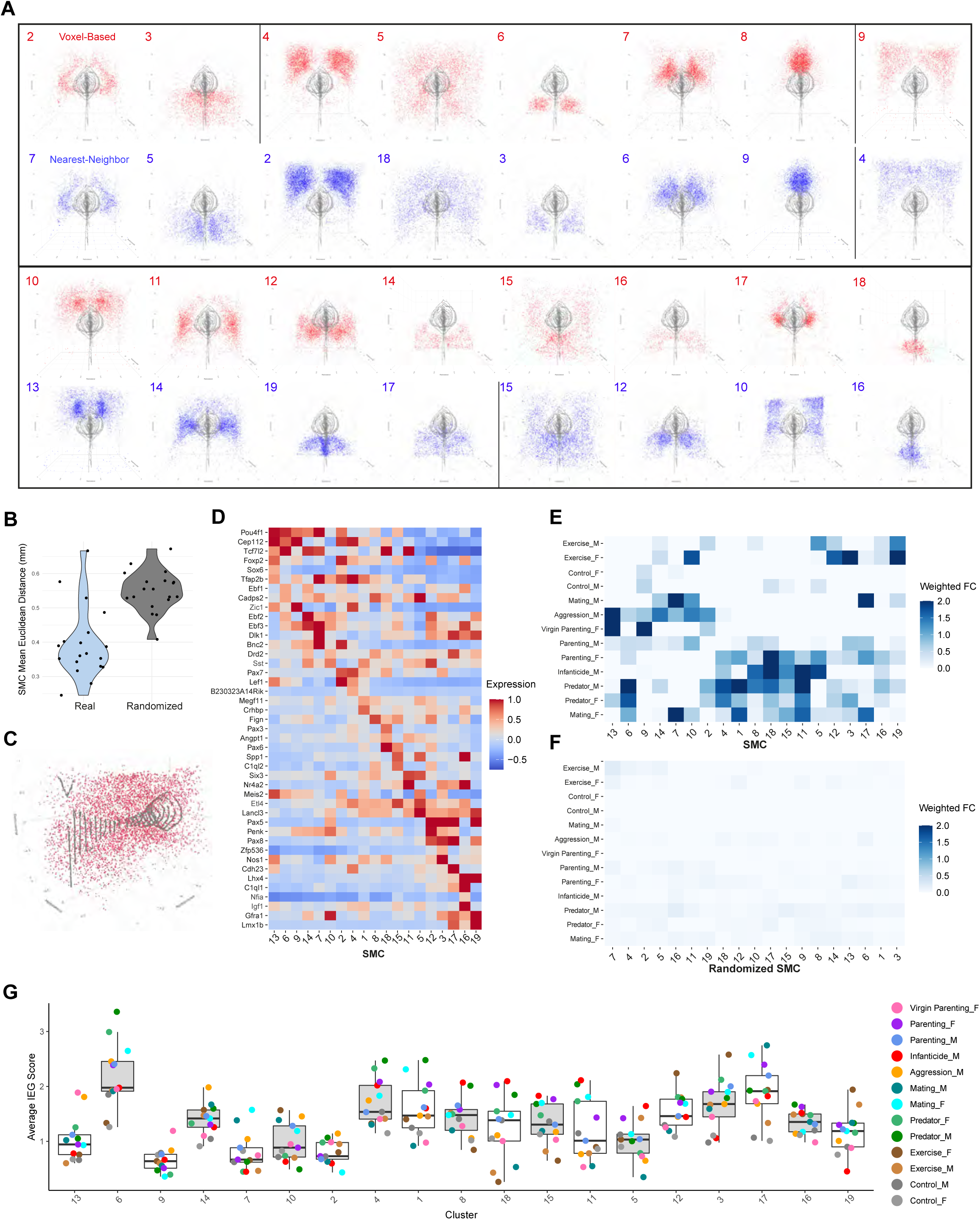
Z-score Expression of Spatially Autocorrelated genes, Related to Figure 3 and Table 2. (A) Comparison of window-defined SMCs (red) and the original SMCs (blue). (B) Violin plot illustrating the compactness of SMCs by calculating the mean distance for all cells in each real and randomized SMC. (C) A representative image of a randomized SMC showing no spatial pattern. (D) Z-score expression of genes demarcating SMCs. (E) Weighted fold change for each behavior in real SMCs and (F) simulated SMCs. (G) IEG Scores for each behavior in real SMCs.

**Figure S5:**
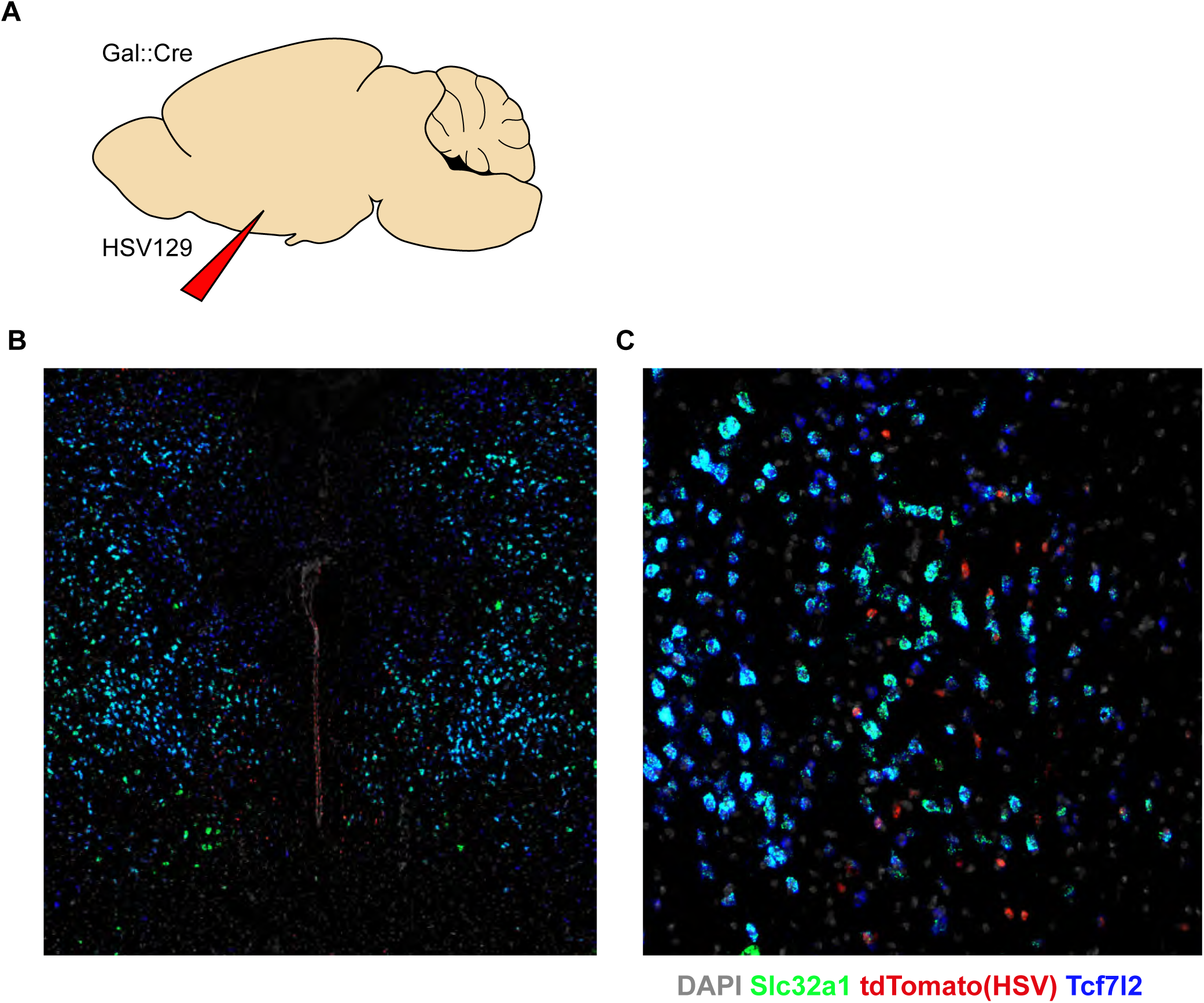
Identification of Presumptive POAGal Inputs to Rostral PAG, Related to Figure 6. (A) Paradigm of conditional HSV129 injection into POA of Galanin::Cre mice. (B) RNAScope in Situ hybridization of rostral PAG 48 hours post injection (grey: DAPI, green: Slc32a1, red: tdTomato, blue: Tcf7l2) (C) Zoomed view of (B) showing overlap of HSV/tdTomato-positive cells with Tcf7l2 and Slc32a1.

**Figure S6:**
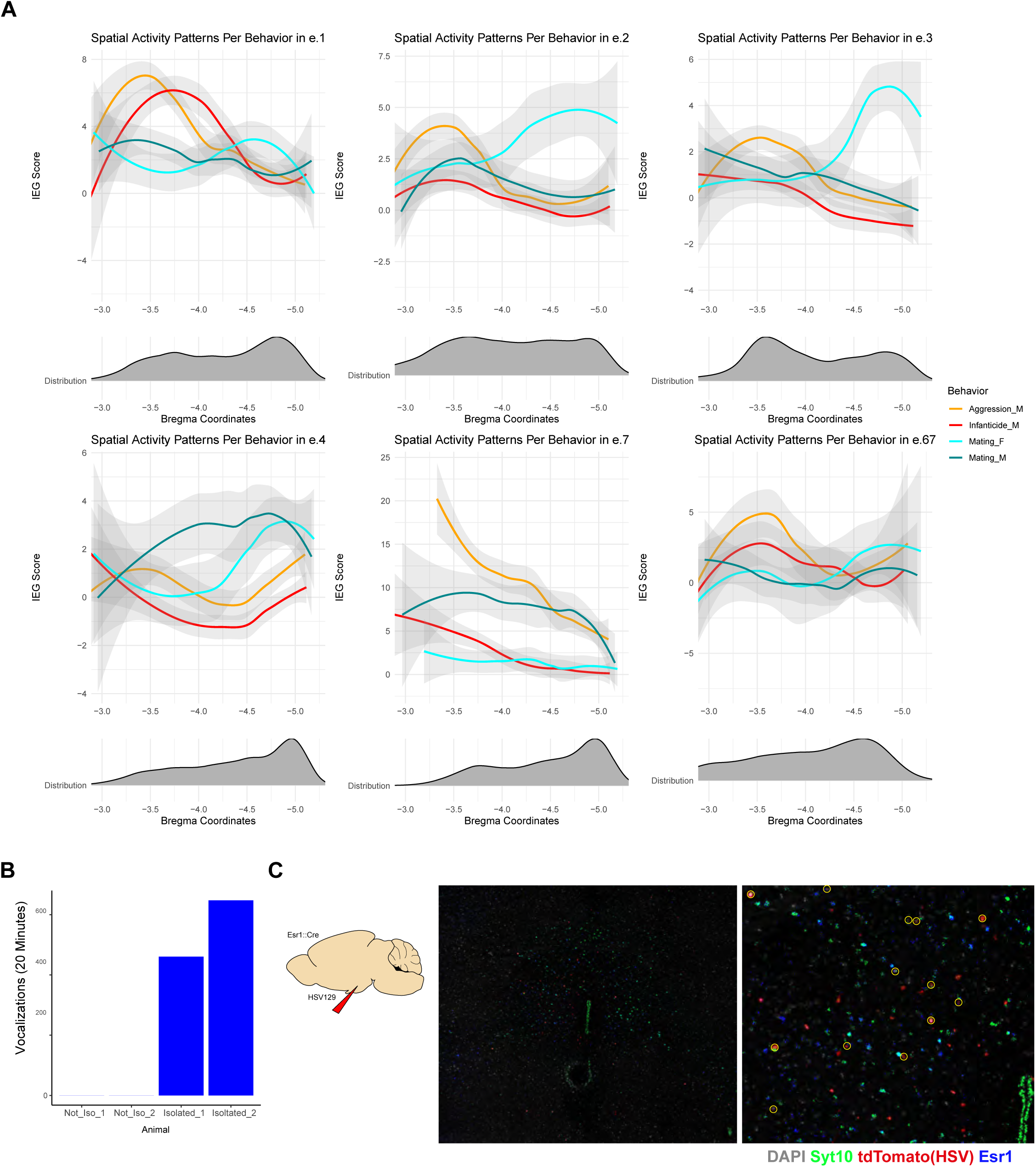
Activation of Esr1+ Clusters Along the Rostrocaudal Axis, Related to Figure 7. (A) (Top) Smoothed, per-behavior IEG Scores along the rostrocaudal axis. (Bottom) Distribution of each cluster along the rostrocaudal axis. (B) Number of vocalizations of four pups after 20 minutes in non-isolated condition (with other pups) and isolated condition. (C) (Left) Paradigm of conditional HSV129 injection into VMHvl of Esr1::Cre mice. (Middle) RNAScope in Situ hybridization of PAG 48 hours post injection (grey: DAPI, green: Sy10, red: tdTomato, blue: Esr1). (Right) Zoomed view of showing overlap of HSV/tdTomato-positive cells with Syt10 and Esr1.

## STAR METHODS

### Animals

Adult C57Bl6/J male and female mice aged 50-90 days were used in this study. Animals were kept on a 12 hour:12 hour light/dark cycle with *ad libitum* food and water. Animal care and experiments were conducted in accordance with NIH guidelines and were approved by the Harvard University Institutional Animal Care and Use Committee (IACUC).

### Single-cell nuclear sequencing and library preparation

Adult virgin male and female mouse brains (aged ∼7-8 weeks) were harvested using a metal brain matrix with 1mm coronal slicing slots. From the level of the thalamus to the cerebellum, three 1mm slices were taken and the periaqueductal gray (PAG) dissected from each. Tissue was immediately placed into ice cold dissociation buffer made of: 10mM NaCl, 3mM MgCl2 and 0.1% NonIdet P40. Tissue was then homogenized using a 3mL Potter-Elvehjem Tissue Grinder at 300rpm within a 4C cold room. The liquid was then transferred to a 15mL tube and centrifuged at a low speed (to reduce stress induced artifacts) of ∼50rcf for 5 minutes at 4C. After decanting supernatant, the pellet was gently resuspended in 1.5mL of 1% BSA PBS with .2U/uL Promega RNasin. Note that higher centrifugation in the previous step will increase nuclei recovered at the cost of a much stronger pellet that will increase trituration time. The suspension was then passed over a 70um filter, followed by a 20um filter all performed on ice. We stained nuclei by adding 1.5uL of DAPI to the suspension and gently mixing and incubating for 5 minutes prior to Fluorescence-activated Cell Sorting (FACS). FACS was used to sort 500,000 events into a 1.5mL Eppendorf tube. From the sorted suspension, 1uL of propidium iodide was added to 10uL suspension to estimate the nuclear concentration with a Luna Cell Counter.

The nuclear suspension was then loaded into a 10X Chromium Single Cell 3’ (v2 chemistry) chip at the appropriate concentration, followed by standard 10X Chromium Single Cell 3’ (v2 chemistry) protocols. The resulting libraries were sequenced on a NovaSeq 6000 S4 200 flowcell to a sequencing saturation of at least 75% for each run.

### Single-nucleus sequencing analysis

Sequencing reads were aligned using 10X Genomics Cellranger 2.1 to the mouse genome (mm10 v1.2) with intronic reads included. The mean number of UMIs and genes per cell were 3,438 UMIs and 1,716 genes for excitatory neurons and 3,093 UMIs and 1,595 genes for inhibitory neurons. Raw reads and output are available at: https://data.nemoarchive.org/biccn/grant/u19_huang/dulac/transcriptome/sncell/10x_v2/mouse/pag/.

For analysis, we relied primarily on the R package Seurat and standard data analysis practices (Macosko et al., 2015). We started our analysis by first filtering out any cell with less than 500 UMIs or more than 40,000 UMIs. Additionally, to remove confounding sources of variation from our clustering, we defined the percentage of counts derived from problematic genes related to mitochondria, ribosomes, red blood cells (RBCs), apoptosis, or immediate early genes (IEG). Cells with more than 10% mitochondrial, 1% ribosomal, 2% RBC, 0.2% IEG, or 0.2% apoptotic genes were filtered out.

As a first pass, we defined major cell classes (Oligodendrocyte, neuron, astrocyte, etc.). Regions that could clearly be identified as non-PAG (via multiple markers and Allen Brain Atlas in situ references) were excluded from clustering analysis on the PAG. To cluster neurons, inhibitory and excitatory neurons were split, with monoaminergic clusters included in the excitatory subset. Variable genes were defined using the Seurat function ‘FindVariableGenes’. To ensure clustering was not influenced by any cellular stress events, variable genes belonging to the problematic genes noted above were removed. Variable genes were then scaled and regressed using the Seurat function ScaleData with nCount_RNA, percent.mito, percent.ribo, age, batch used as the variables to regress and the ‘negbinom’ model used. These variable genes were used to define 80 principal components that were then used to build a shared nearest neighbor network (SNN) with k = 15. The leiden algorithm was used to cluster neurons with resolution set to 2.4. To define stable clusters, we assessed several clustering resolutions and utilized a bootstrapping approach where 80% of the cells were randomly chosen to define clusters. Pairwise Jaccard coefficients were then calculated for each bootstrap subset of cells compared to the clustering performed at the same resolution with 100% of cells. Clusters were deemed stable if the Jaccard coefficient was greater than 0.4. The resolution that provided the highest number of stable clusters was chosen. Cells from unstable clusters were merged back into the data by training a support vector machine classifier on stable clusters and assigning each unstable cell to a cluster. The same approach was applied to inhibitory neurons but with k=11 and resolution set to 2.4.

### Selection of MERFISH Gene Panel

To identify transcriptionally distinct cell populations with MERFISH, we designed a panel of 262 marker genes. This panel included 12 genes that were manually selected because they are well-characterized markers of a major cell type, with at least one such marker for neurons, inhibitory neurons, excitatory neurons, mature glial cells, astrocytes, mature oligodendrocytes, oligodendrocyte precursor cells, microglia, macrophages, endothelial cells, ependymal cells, and mural cells. The remaining genes were computationally selected to discriminate between the set of neuronal clusters identified with snRNA-Seq using two approaches. We first calculated markers for excitatory and inhibitory neurons using the Seurat function FindAllMarkers with test.use = ‘negbinon’. We then filtered for genes that 1) were expressed in less than 40% of cells and 2) exhibited a bonferroni corrected p-value <0.01. We then used a minimum-redundancy maximum-relevancy algorithm (MRMR) to identify a set of 150 genes that exhibited the highest mutual information with the snRNA-Seq clusters while also exhibiting the lowest mutual information with the other selected genes (De Jay et al., 2013). Put another way, this approach seeks to select genes with high information and remove genes with redundant information (i.e. if three genes are equally good markers for cluster 1, then select only one). Because this approach tends to bias for highly expressed genes, we additionally selected ∼40 other biologically informative differentially expressed genes not included in the MRMR set. To further improve this gene set, we trained a radial support vector machine classifier (5-fold repeated cross validation) using the 200 genes on 50% of the snRNA-Seq neurons. We then classified the left-out neurons and iteratively replaced genes to boost poor accuracy clusters while also utilizing genes with more binary expression (on or off as opposed to graded expression).

The second approach used differential expression analysis between all pairwise comparisons of the snRNA-Seq clusters, as described previously (Zhang et al., 2020). Briefly, for a given pair of clusters, the subset of genes that were 1) at least 2-fold higher in expression in cluster A relative to cluster B, 2) expressed in at least 10% of cluster A cells, 3) expressed in at least 3-fold greater a fraction of cluster A cells than cluster B cells, and 4) significantly greater in expression in cluster A relative to cluster B based on having a p-value < 0.05 in a t-test. This set of genes was then sorted based on their p-value, restricted to no more than the top 50 genes, and assigned a score from 10 to 1 based on its position in the list (i.e. the genes with the 5 lowest p values were scored 10, the next 5 lowest were scored 9, etc.). This process was repeated for all possible combinations of clusters. The total score for each gene was computed as the sum of its scores from all the lists, and the gene with the highest score was selected. The highest scoring gene was selected until there were at least two genes from a given comparison present in the selected set, at which point that list of genes from that comparison was no longer used in calculating the score of the genes. This process was repeated until there were at least 2 differentially expressed genes from all pairwise comparisons. We performed the differential expression analysis independently on the inhibitory clusters and the excitatory clusters and selected 65 genes for inhibitory clusters and 64 genes for excitatory clusters, with the union of these lists yielding a total of 116 unique genes. Combining the lists of genes selected by hand, based on MRMR, and based on differential expression yielded 262 unique genes. Four of these genes were relatively highly expressed and were thus selected to be imaged individually in their own rounds of sequential FISH imaging, with the remaining 258 genes imaged with combinatorial FISH.

### Design and construction of MERFISH encoding probes

MERFISH encoding probes for the 258 combinatorial marker genes were designed as described previously. Starting with a codebook of 285 unique binary barcodes constructed from 20 bits and designed to each have a Hamming weight of 4 and a Hamming distance of at least 4, each of the 258 genes was paired with a barcode, and the remaining 27 unassigned barcodes were left as “blanks” (i.e. barcodes not assigned to any gene that provide an estimate of the false-positive rate in MERFISH).

The same procedure was used for the 26 activity-related genes, except using a codebook constructed from 10 bits that had 30 unique barcodes with these parameters. The barcode of a gene determines the readout sequences that will be associated with its encoding probes.

All possible 30-mer targetable regions for the marker genes were designed as described previously. Briefly, all 30 nt regions with a GC fraction between 0.4 and 0.6 and a melting temperature between 60 and 75 C were identified, and the set of them that overlapped by no more than 20 nt were selected as potential target sites. To construct the encoding probes, 64 unique target regions were selected at random from the set of potential target sites on a given gene, one readout sequence was prepended to this sequence, and two additional readout sequences were appended to the sequence, separating each portion with an A. In all cases the three readout sequences associated with a given targeting sequence were different from eachother, and all possible combinations of readout sequences and orders were used with equal frequency for a given genes encoding probes when possible. A subset of genes had fewer than 64 potential target sites, in this cases they were targeted with as many probes as possible, resulting in Arpp21 targeted with 25 probes, Otx2os1 targeted with 58 probes, Rln3 targeted with 39 probes, Sept4 targeted with 33 probes, Cck targeted with 52 probes, Sst targeted with 49 probes, Sec61b targeted with 39 probes, Prlr targeted with 44 probes, Npy targeted with 48 probes, Lmo1 targeted with 25 probes, Npw targeted with 43 probes, and Pth2 targeted with 39 probes. These genes may have a lower signal-to-noise ratio than those targeted by 64 probes. To create the final sequences, we placed a forward and reverse primer on the 5’ and 3’ end of the encoding probe, respectively. These sequences were ordered as an oligo pool (Twist biosciences), and amplified and purified as previously described. The final complex pool of encoding probes was further purified on a 4% urea-polyacrylamide gel. The same procedure was used to construct the encoding probes for the activity-related genes, and in this case all genes of interest were targeted with 64 encoding probes. The marker genes imaged in sequential rounds were assigned probes as described above, but in this case they were each targeted with 48 probes, and a single readout sequence was assigned to each gene and was present in two copies on each encoding probe for that gene.

### Staining tissue with MERFISH encoding probes

Samples were stained for MERFISH imaging as previously described. Briefly, after a coverslip containing a set of brain slices had incubated at 4 C in 70% ethanol overnight, it was brought to room temperature and washed twice with 2x saline sodium citrate (2x SSC). The coverslip was then washed once with 30% formamide wash buffer (30% formamide, 2X SSC) for 5 minutes at room temperature. A hybridization chamber was prepared for the coverslip by placing parafilm onto a petri dish, and adding a 100 µL droplet of encoding probe hybridization mixture. Encoding probe hybridization mixture was prepared by taking 100 µL of hybridization buffer (30% formamide, 1 mg/mL yeast tRNA (Life Technologies, 15401-011), 10% v/v dextran sulfate (Sigma, D8906), 2x SSC) and adding the encoding probe pools of combinatorial marker genes and activity-related genes to a final concentration of 0.1 nM per probe, the pool of sequential marker genes to a final concentration of 0.9 nM per probe, and the poly(A) anchor probe (composed of DNA and LNA nucleotides TTGAGTGGATGGAGTGTAATT+TT+TT+TT+TT+TT+TT+TT+TT+TT+T, were T+ is an LNA, the probe has an acrydite at the 5’ end that covalently links it to a polyacrylamide gel, below, hybridizes to the poly(A) tail of mRNAs thus linking them to the polyacrylamide gel, and has a unique readout sequence) to a final concentration of 1 µM. The coverslip was inverted onto the droplet, and the coverslip was incubated in a humidified incubator at 37 C for 36 to 48 hours in the covered petri dish.

After hybridization the sample was embedded in a polyacrylamide gel and cleared as described previously. Briefly, the sample was washed twice at 47 C with 30% formamide wash buffer for 30 min per wash. The sample was washed in 2x SSC for 5 min, then washed twice in 4% polyacrylamide gel mixture (4 % acrylamide [19:1 with bis], 50 mM Tris-Hcl, pH 8, 300 mM NaCl, 0.05% v/v Temed, 0.05% wt/v Ammonium persulfate, degassed), then inverted onto a droplet of 4% polyacrylamide gel mixture and incubated at room temperature for 2 hours until the gel was polymerized. The embedded sample was cleared with an SDS proteinase K digestion buffer (2% v/v sodium dodecyl sulfate (SDS; ThermoFisher, AM9823), 0.5% v/v Triton X-100 (Sigma, X100), and 1% v/v proteinase K (New England Biolabs, P8107S), and 2×SSC) for 48-72 hours at 37 °C. After digestion, the coverslips were washed four times with 2x SSC for 30 minutes per wash, then stored at 4°C in 2x SSC with 1:100 Murine RNase inhibitor (New England Biolabs, M0314S) prior to imaging.

### MERFISH imaging

Imaging was performing on a custom-built imaging system that we described previously (Moffitt et al., 2018). Prior to loading the coverslip onto the microscope, the sample was stained with the readout probe complementary to the readout sequence present on the poly(A)-anchor probe (readout probe was conjugated via a disulfide bond to Alexa488 and stained in hybridization and wash buffer at a final concentration of 3 nM) for 15 min at room temperature, followed by staining with DAPI (1 µg/mL in hybridization and wash buffer) for 10 min at room temperature, after which the sample was placed in 2x SSC and loaded onto the microscope. Hybridization and wash buffer is 2x SSC, 10% v/v ethylene carbonate (Sigma, E26258), and 0.1 % v/v Triton X-100, and when used for readout probe hybridization is supplemented with the appropriate readout probe(s) to a final concentration of 3 nM each. The coverslip was placed into a commercial flow chamber (Bioptechs, FCS2), with a 0.75 mm thick silicone gasket to provide space for liquid flow. Imaging buffer was flowed into the chamber (5 mM 3,4- dihydroxybenzoic acid (Sigma, P5630), 2 mM trolox (Sigma, 238813), 50 µM trolox quinone, 1:500 recombinant protocatechuate 3,4-dioxygenase (rPCO; OYC Americas), 1:500 Murine RNase inhibitor, and 5 mM NaOH (to restore pH to 7) in 2x SSC) and an initial scan of the coverslip was performed at low magnification (Nikon, CFI Plan Apo Lambda 10x) with 405-nm illumination to produce a low-resolution mosaic of the DAPI signal in all slices. Based on the mosaic image, the approximate center of the cerebral aqueduct was identified in each slice and a 9 by 9 grid of field-of-view (FOV) positions was generated, with the center of each FOV separated by 200 µm. Automated imaging was then performed with a high magnification, high-numerical aperture objective (Olympus, Super Apochromat 60x with 1.3 NA) at each FOV position. The first round of imaging was performed with a light path set through a spinning disc confocal (Andor Yokogawa CSU-W), collecting images in the 560 nm, 488 nm, and 405 nm channels to image the fiducial beads, the poly(A) RNA stain, and DAPI, respectively. A single z-plane at the surface of the coverglass (z = 0 µm) was acquired for 560 nm channel, and 6 z-planes were acquired for the 488 nm and 405 nm channels (z = 0, 1.5, 3, 4.5, 6, 7.5 µm). After the first round of imaging, the disulfide bond on the readout probes were cleaved flowing 2.5 mL of cleavage buffer (100 mM Tris (2-carboxyethyl) phosphine (TCEP; Sigma, 646547) in 2x SSC) and incubating for 9 min in the flow chamber at room temperature. The sample was washed with 1.5 mL of 2x SSC, and then the hybridization of the readout probes for the next round were performed. Hybridization was performed using the hybridization and wash buffer described above containing the appropriate readouts, 3.5 mL was flowed across the chamber and the sample was incubated for 18 minutes. The sample was then washed with 1.5 mL of hybridization and wash buffer that contained no readout probes for 15 minutes. Finally, 1.5 mL of imaging buffer was flowed onto the sample and the next round of imaging was performed. Following the first round of imaging, the light path was shifted to acquire widefield images, and in all cases the 750-nm, 650-nm, and 560-nm channels were imaged to visualize the 2 readout probes and fiducial beads. When a readout probe corresponded to a bit being used for a combinatorial marker gene we acquired 6 z-planes (z = 0, 1.5, 3, 4.5, 6, 7.5 µm), otherwise we acquired a single central z-plane (z = 4.5 µm), and beads were always acquired at z = 0 µm. The cycle of cleavage-hybridization-imaging was repeated until all the bits in the combinatorial marker gene encoding probes, combinatorial activity-related gene probes, and sequential marker gene probes were imaged. A final round of imaging was performed to acquire a background measurement for the sequential marker genes by cleaving the probes and then flowing on imaging buffer as before, without performing a hybridization. Imaging of the blank round was performed for the 750-nm, 650-nm, and 560-nm channels at z = 4.5 µm, z = 4.5 µm, and z = 0 µm, respectively. In total 19 rounds of imaging were performed, with 17 rounds of imaging to visualize the bits used to encode all the genes, one round to visualize cell nuclei and cytoplasmic extent, and one round to visualize background. All buffers and readout probe mixtures were flowed with a custom built, computer-controlled fluidics system composed of four, 8-port valves (Hamilton, HVXM 8-5) and a peristaltic pump (Rainin, Dynamax RP-1), configured as previously described (Chen et al., 2015).

### MERFISH image analysis and cell segmentation

All MERFISH image analysis was performed using MERlin (Emanuel, 2020) (https://github.com/emanuega/MERlin), a Python-based MERFISH analysis pipeline, using algorithms similar to what we have described previously (Moffitt et al., 2016; Xia et al., 2019). Analysis begins by aligning the images from each imaging round based on the fiducial bead images, thus correcting for X-Y drift in the stage position that may occur over the course of the experiment. Images were then high-pass filtered to remove background, RNA spots were tightened with 20 rounds of Lucy-Richardson deconvolution to better resolve closely positioned spots, and then low-pass filtered to partially re-spread the spots and thereby help account for any slight mis-alignment between rounds. To identify RNA molecules we used the pixel-based decoding algorithm we previously described (Moffitt et al., 2016) and filtered the resulting putative RNA molecules based on several properties (i.e., overall intensity, similarity to assigned barcode, and number of pixels contributing to the molecule) as previously described (Xia et al., 2019) to obtain a gross barcode misidentification rate of 10%. The decoding and filtering was performed independently for the two combinatorial gene sets, but with all images aligned together such that they had identical coordinate systems. Cell boundaries were identified using a seeded watershed algorithm applied as previously described (Moffitt et al., 2018), using the DAPI images to identify seeds and the poly(A) RNA stain to identify the cytoplasmic extent. RNA molecules were partitioned into cells if they fell within the cell boundary, yielding a counts matrix for each cell. For the genes measured with sequential two-color FISH, the signal from these imaging rounds was quantified by summing the fluorescence intensity of all pixels that fell within the segmentation boundary of a cell in the same z-plane as was acquired for the image and then normalized by the number of pixels that contributed to the signal. The signal was further corrected based on the channel-specific background measurement for the cell. Cases where a cell lacked a boundary in the z plane acquired for the sequential measurements were assigned as nan. These signals are included in the counts matrix.

### MERFISH Clustering Analysis

The segmented, normalized cell-count matrix output from MERlin was loaded into Seurat for further processing. We first clustered cells into major cell classes (neurons, glia, smooth muscle cells, etc.). As our data intentionally included neuron-centric genes, we did not focus our analysis on glial subdivisions that may exist but are not evident from the minimal gene set used to identify them. Upon parsing out neurons, we filtered cells with high amounts of glial-specific genes or cells with less than 40 counts detected in the cell. We further subset excitatory and inhibitory neurons, further separating excitatory neurons into excitatory neurons within the PAG, and other highly discrete nuclei/structures surrounding the PAG: colliculi, cerebellum, dorsal raphe, occulmotor, Edinger Westphal and trochlear, laterodorsal tegmental nuclei. For all neurons, we calculated z-score on each replicate to reduce technical variation. For inhibitory and excitatory neurons, we defined 50 and 60 principal components (PCs), respectively, using genes expressed in neurons. We used these PCs to build a shared nearest neighbor network with k=11, followed by Louvain clustering with resolution set to 2.6 and 3 for inhibitory and excitatory neurons, respectively. To define stable clusters, we used same approach applied to nuclear sequencing data specified above.

### Calculation of Immediate Early Gene (IEG) Score

We sought to use multiple IEGs to help compensate for the noise inherent to any gene’s expression. From a list of 26 IEGs listed in Table 1, we further refined this list down to 7 (Fos, Egr1, Fosl2, Fosb, Jun, Nr4a1, Nr4a3) that exhibited correlation with each other, while others had poor signal or didn’t display activity dependent expression. To reduce the effect of small batch effects from replicate to replicate that result in slightly shifted non-zero distributions, we sought to remove these batch effects by normalizing the IEG distributions across all replicates. To do this, we calculated the median minimum and maximum of each IEG’s non-zero distribution across all replicates and used these values to rescale the non-zero distribution for each gene, thus, giving each IEG a common floor and ceiling of expression. Normalized IEG values were further z-scored by replicate. To calculate IEG Scores, we summated the z-scores of the seven genes. For weighted fold-change, we divided the cluster’s mean IEG Score for a given behavior by the mean IEG Score for all behaviors in that cluster, minus 1. This was followed by the addition of the difference between mean IEG Score for a given behavior and the cluster mean IEG Score for all behaviors.

#### Statistical Tests for Activations

We performed unpaired Student’s t-tests for each cluster by comparing the IEG Scores (for all cells in that cluster) of each behavior to that of all other behaviors. After, p-values were corrected for multiple comparison’s using the false discovery (fdr) metric. Clusters were deemed significantly activated if fdr ≤ 0.05, the difference in behavior-IEG Score and average cluster IEG Score > 0.3, and the given behavior-IEG Score > 1. In this test, significant clusters represent more selective activation above that of all other behaviors. However, we also sought to compare cluster activities to that of naïve animals alone. Accordingly, we separately performed unpaired Student’s t-tests (with fdr correction) on each cluster, comparing IEG Scores for each behavior to that of naïve male and female cells belonging to the same cluster. This test was aimed at identifying if a cluster is activated during a behavior at all, regardless of whether it is also activated in every other behavior or only one. Hence, we noted clusters identified as activated compared to naïve animals are more likely to match the appearance of Fos RNAScope in situ hybridizations, while the comparison to all other behaviors is more likely to connote increased relevance for a specific behavior(s).

### Calculation of Spatial Networks and Spatial Autocorrelation

#### Definition of coordinates

Because MERFISH slices vary from batch to batch and slice to slice, we sought to effectively standardize their spatial attributes by 1) defining the center of the aqueduct by taking the median x and y coordinates of ependymal cells, then 2) defining each neuron’s distance from this center by subtracting their x and y coordinates from the slice-defined aqueduct center. Anterior-Posterior (AP) Bregma coordinates for each replicate were estimated by examining the first slice from each replicate and manually defining which slice is the most anterior. This slice was defined as AP: -2.7mm by comparison to the Paxinos mouse coordinate system. After the starting point was defined, we could then align all other replicates by aligning their distributions of spatially restricted clusters. To do this, we selected six spatially localized clusters to align them by. Per replicate, we defined each cluster’s mean z position, zµ. We then calculated the mean zµ for each cluster across all replicates, clz. We then defined the relative AP shift for each replicate by, for all six clusters, calculating the mean of clz - zµ, Δz.

Effectively, Δz then reflects how many slices (100um distance from each) a given replicate is shifted relative to another, i.e. for any slice, z, a replicate with Δz = -1.2 is approximately 120um more anterior than a replicate with Δz=0. These values were used to assign AP positions for each replicate, assuming a starting point of -2.7mm for the replicate with the minimum Δz.

#### Nearest-Neighbor Network Construction and Spatial Metacluster Definition

We then used the three-dimensional coordinates defined above to, independently for each replicate, build a k =25 nearest neighbor network using the RANN R package (https://cran.r-project.org/web/packages/RANN/index.html). We used this graph to build an adjacency matrix and subsequently tallied the number of neighbors between each cluster, with self-connections removed. We hierarchically clustered the resulting matrix using the R function hclust with the method = ‘ward.D’. The object was then fed into a dynamic treecut algorithm (Langfelder et al., 2008) with variables set to: minClusterSize = 1, method = ’hybrid’, deepSplit = 4. To confirm our approach was not creating seemingly spatial patterns in potentially random data, we repeated this process with randomized connections by shuffling the column/cell names of the adjacency matrix. This resulted in five, randomly distributed metaclusters with no apparent spatial patterns (data not shown).

#### Moran’s I Calculation

For the calculation of spatial autocorrelation in Table 2 we utilized Moran’s i statistic. A k=15 nearest neighbor graph was constructed, as mentioned above, on the most thoroughly sampled replicate, Aggression Male 2, which included 25 slices separated by 100um. We converted the object to a nearest-neighbor list using the nb2listw function in the spdep R package (Bivand and Wong, 2018). For each gene’s z-score expression, we then used this list to perform a permutation test for Moran’s I statistic calculated using 100 random permutations of a given gene expression vector for the given spatial weighting scheme.

### Behavioral Assays

Animals were individually housed for ∼3-4 days before behavioral assays, which were performed as described previously (Moffitt et al., 2018; Wu et al., 2014), beginning >1 hour after the onset of the dark cycle under dim red lighting. Mice were habituated to the testing room for 1 hour before the start of each behavioral assay. Only mice that performed the desired behaviors (described below) after the defined behavioral stimuli were selected for tissue collection.

#### Parental behavior

Virgin females: A single mouse pup aged 2-3 days was introduced into one corner of the home cage of an adult virgin female mouse (age ∼8 weeks), away from the nesting material. A parental response was defined as retrieval of the pup to the nest combined with nesting, crouching, and grooming. Mice were sacrificed for tissue harvesting 30 minutes following the retrieval of the pup to the nest. Only mice displaying all the above behavioral subroutines were selected for further processing.

Mothers and Fathers: Virgin male and female mice (age ∼7 weeks) were co-housed in pairs until the birth of pups (∼21 days). Mice were allowed to remain with pups for 3 days before mothers were moved to a fresh cage and pups were removed from the home cage of the father. Two days later, parental behavior was tested as described above with virgin females.

#### Pup-directed aggression

A single mouse pup aged 2-3 days was introduced into one corner of the home cage of an adult virgin male mouse (age ∼8 weeks), away from the nesting material. Mice were sacrificed for tissue harvesting 30 minutes following aggressive attacking of the pup, characterized by biting behavior leading to audible pup vocalizations and occasionally preceded by tail rattling. Pups were immediately removed from the cage following an attack.

#### Inter-male aggression

C57Bl6/J juvenile males (∼4 weeks) were introduced into the home cage of an adult virgin male mouse (age ∼8 weeks). Mice were sacrificed for tissue harvesting 30 minutes following aggressive attacking of the intruder, characterized by rapid bouts of biting leading to defensive reactions by the intruder (rearing and escaping).

#### Male mating

Receptive female mice (∼8 weeks) were placed into a male’s cage which was followed by mating. Mice were sacrificed for tissue harvesting 30 minutes following ejaculation. Mounting intromission bouts usually occurred 4-6 times before males ejaculated. The female mouse was left in the cage following ejaculation. Males exhibited decreased activity and generally sat in their nest without interaction post-ejaculation.

#### Female mating

An intact adult sexually experienced male mouse was introduced into the home cage of an adult virgin female mouse (age ∼8 weeks) in estrous (determined by a vaginal smear). Females were sacrificed for tissue harvesting 30 minutes following intromission by the male. Males were removed from the cage 10 minutes after the first intromission.

#### Toy Snake Fear

Male and female mice (∼8 weeks) were placed in a 2x2 ft. open field container with two red huts placed centrally, approximately 3 inches from the wall. Mice were habituated to the open field for an hour pre- snake exposure and explored all sides normally, crossing through the middle of the field often. Three realistic toy snakes were then placed down the midline of the open field, with one snake manually moved from the head using fishing line for ten minutes. Mice robustly exhibited wall rearing in the corners and occasional outstretched investigations of the snakes before darting back to a red hut or corner. While mice initially urinated or defecated little to none during the habituation period, they defecated and urinated several times after the introduction of snakes. After ten minutes of snake movement, the snakes were left motionless down the midline of the open field, before being carefully removed. At no point during snake exposure did mice touch the snake or cross the midline, indicating a strong negative valence. Brains were harvested 30 minutes after the ten-minute snake movement period.

#### Voluntary Wheel Exercise

Male and female mice (∼8 weeks) were habituated to wheels for one week prior to testing (3 hours of wheel exposure in testing room every other day). On the testing day, wheels were introduced to the home cages of mice, where they readily ran nearly the entire time after wheel introduction. After an hour of almost constant wheel running, brains were harvested from mice.

### Stereotaxic Viral Injections

All surgeries were performed under aseptic conditions in animals anaesthetized initially with isoflurane followed by injection of 100 mg kg−1 ketamine (KetaVed, Vedco) and 10 mg kg−1 xylazine (AnaSed) via intraperitoneal (i.p.) injection.

For anterograde tracing, 100nl of HSV1-H129ΔTK-TT virus was injected into the VMHvl of Esr1-Cre, MPOA of Gal-Cre, or AvPe of Ucn3-Cre mice, under BL2 safety guidelines using a Nanoject III injector (Drummond Scientific). Mice were monitored for overall health and sacrificed 48-60 hours later. For monosynaptic rabies tracing, a 1:1 ratio (150 nL) of AAV1-EF1a-FLEX-TVA-mCherry (avian TVA receptor, UNC Viral Vector Core) and AAV1-CAG-FLEX-oG-WPRE-SV40-PA (optimized rabies glycoprotein, Salk Institute) was stereotaxically injected into the PAG of Esr1-Cre males and females. For all tissue collection, mice were transcardially perfused with 4% paraformaldehyde in PBS, followed by overnight post-fixation in PFA and cryoprotected with 30% sucrose.

### In situ hybridization

Double and triple label fluorescent in situ hybridization was performed using the RNAscope assay V2 kit (Advanced Cell Diagnostics, ACD) according to the manufacturer’s instructions. Frozen brains were sectioned on a cryostat at 16Dm and stored at -80° C. Slides were thawed and fixed in 4% PFA for 15 minutes followed by dehydration in 50%, then 75%, then 100% ethanol. Cells were permeabilized for 30 minutes at room temperature using Protease III provided by ACD. Sections were then processed as suggested by the ACD protocol and mounted with DAPI-included mounting media. Slides were imaged at 10x or 20x on an Axioimager (Zeiss). All probes were made by ACD.

